# Single-cell sequencing of human iPSC-derived cerebellar organoids shows recapitulation of cerebellar development

**DOI:** 10.1101/2020.07.01.182196

**Authors:** Samuel Nayler, Devika Agarwal, Fabiola Curion, Rory Bowden, Esther B.E. Becker

## Abstract

Current protocols for producing cerebellar neurons from human pluripotent stem cells (hPSCs) are reliant on animal co-culture and mostly exist as monolayers, which have limited capability to recapitulate the complex arrangement of the brain. We developed a method to differentiate hPSCs into cerebellar organoids that display hallmarks of *in vivo* cerebellar development. Single-cell profiling followed by comparison to an atlas of the developing murine cerebellum revealed transcriptionally-discrete populations encompassing all major cerebellar cell types. Matrigel encapsulation altered organoid growth dynamics, resulting in differential regulation of cell cycle, migration and cell-death pathways. However, this was at the expense of reproducibility. Furthermore, we showed the contribution of basement membrane signalling to both cellular composition of the organoids and developmentally-relevant gene expression programmes. This model system has exciting implications for studying cerebellar development and disease most notably by providing xeno-free conditions, representing a more biologically relevant and therapeutically tractable culture setting.

## INTRODUCTION

The cerebellum has a major role in controlling sensorimotor functions but is also increasingly implicated in higher-order cognitive functions including language, emotion and reward. As a consequence, dysfunction of the cerebellum is linked to an increasing number of motor diseases such as ataxia, dystonia and tremor, and non-motor disorders including autism spectrum disorder (Reeber et al., 2013). Specifically, abnormal cerebellar development is an emerging theme contributing to many brain disorders (Sathyanesan et al., 2019). The human cerebellum is one of the first structures of the brain to differentiate and continues to develop until several years after birth. This protracted development after birth makes the cerebellum especially vulnerable to insult. However, the molecular mechanisms underlying physiological development of the human cerebellum and how these go awry in developmental disorders remain incompletely understood. The cerebellum follows a highly stereotyped developmental programme to form a well-organized laminar structure, containing one of the largest and most metabolically active neuronal types (the Purkinje neuron), as well as the most abundant neuron in the brain (the granule cell) (Manto et al., 2012). During early development, the isthmic organizer at the mid-hindbrain (MHB) boundary in the dorsal region of rhombomere 1 induces the formation of the cerebellum through the expression of the lineage-specific transcription factors OTX2, GBX2, EN1/2 and PAX2 (Butts et al., 2014; van Essen et al., 2020). Unique to cerebellar development, precursors arise from two distinct germinal zones in rhombomere 1; the ventricular zone (VZ) gives rise to GABAergic neurons (Purkinje cells (PCs), interneurons, GABAergic deep cerebellar nuclei (DCN) neurons), whereas all glutamatergic neurons (granule cells (GCs), unipolar brush cells, glutamatergic DCN neurons) are generated at the rhombic lip (RL). The fourth ventricle, roof plate (RP) and subsequent hindbrain choroid plexus (CP) are responsible for secretion of growth factors required for development of adjacent regions of the cerebellum, including bone morphogenetic proteins (BMPs) to direct RL formation and Sonic hedgehog (SHH) for VZ precursor differentiation (Manto et al., 2012).

Regulatory mechanisms driving cell-fate specification have been studied in various animal models; however, observing cerebellar development in humans poses several significant challenges, including acquisition of relevant material to study. Human induced pluripotent stem cells (hiPSCs) offer a powerful system to model human cerebellar development and its associated disorders. Different methodologies have recently been developed to differentiate hPSCs into cerebellar neurons through the addition of growth factors that reproduce early patterning events *in vitro*, recapitulating MHB barrier establishment and subsequent formation of polarized cerebellar tissue characteristic of RL/VZ expansion (Ishida et al., 2016; Muguruma et al., 2015; Sundberg et al., 2018; Watson et al., 2018). Current models have mainly focussed on the differentiation of hPSC-derived Purkinje cells through co-culture with mouse cerebellar progenitors. Here, we describe a reproducible method to generate hPSC-derived self-organising three-dimensional organoids without the need of mouse co-culture. Single-cell RNA sequencing (scRNA-seq) revealed transcriptionally distinct populations representative of the majority of cerebellar neuronal cell types, including RL precursors, GCPs, GCs, Bergmann glia, glutamatergic DCN, CP, RP and ciliated cells, whose identity and maturity were confirmed through comparison with published murine single-cell data (Carter et al., 2018). As a proof of principle, we show perturbation analysis of the organoids, in this case treatment by Matrigel embedding, designed to recapitulate basement membrane signalling. Global and population-level differential expression analysis yielded important information about organoid composition, maturity and stimuli response.

## RESULTS

### Embryoid body differentiation recapitulates the formation of the isthmic organizer followed by rhombic lip/ventricular-zone expansion

Re-aggregated hiPSCs were treated with a combination of fibroblast growth factor 2 (FGF2) and Insulin to mimic the self-inductive properties of the isthmic organizer and reproduce MHB development (Figure 1A) (Muguruma et al., 2015; Watson et al., 2018). We refer to cellular aggregates as embryoid bodies (EBs) until the appearance of polarized neuroepithelial tissue and confirmation of MHB identity at day 21, from which point they are referred to as organoids. Successful acquisition of MHB fate acquisition was assessed by measuring expression of lineage-specific transcription factors. At day 21, induction of the MHB markers *EN1* (7.66 ± 5.05-fold-change, p-value 0.0346), *EN2* (318.91 ±183.45, p-value 0.0087), and *GBX2* (8.25 ± 34.71, p-value 0.1147) was observed, at the expense of the anterior marker *OTX2* (0.47 ± 73.66, p-value 0.6429) (Figure 1B). Robust MHB acquisition was further confirmed by immunostaining; at day 21 58.31 ± 12.98% cells were positive for EN1 and 40.52 ± 14.31% for GBX2 (Figures 1A,C). At day 35, acquisition of VZ/RL identity was confirmed by measurement of the VZ markers *OLIG2* (17.74 ±7.09 fold-change, p-value 0.0023), *PTF1A* (9.96 ± 5.36, p-value 0.0072), and *KIRREL2* (6.43 ±1.91, p-value 0.4211), and the RL marker *ATOH1* (28.53 ±19.39, p-value 0.0004) (Figure 1D). Consistently, immunostaining of cryosectioned organoids at day 27, 35 and 42 revealed the emergence of extensive pockets of polarized neuroepithelium expressing KIRREL2, one of the earliest markers of VZ precursors (Supplementary Figure S1). On day 60, organoids exhibited robust expression of the neuronal markers TUJ1 and Calbindin, indicating the maturation of VZ-derived neurons (Figure 1A). The majority of Calbindin-positive cells exhibited bipolar morphology consistent with maturing PC progenitors (data not shown).

**Figure 1.**
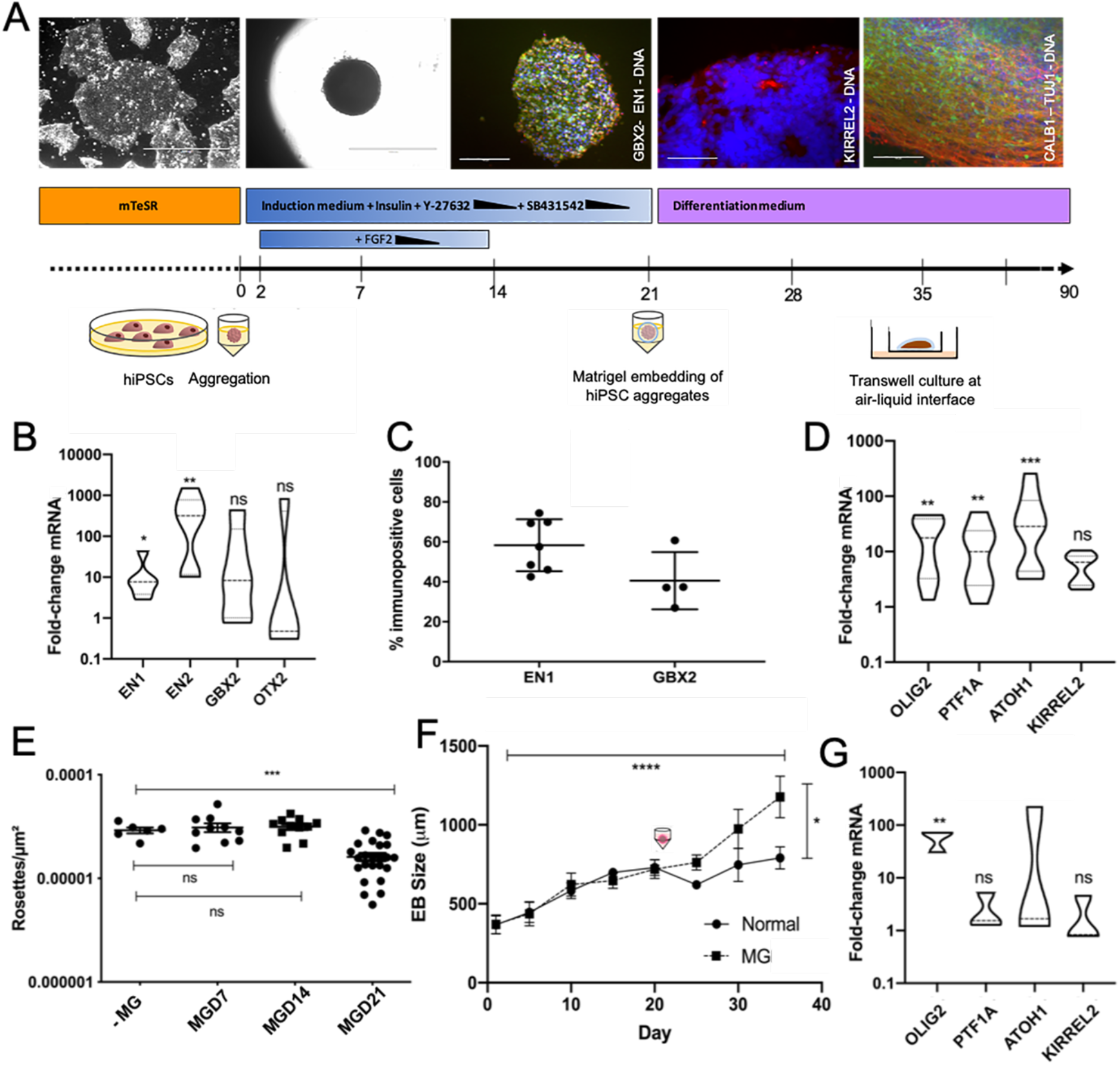
Cerebellar differentiation protocol recapitulates MHB formation and subsequent RL/VZ expansion. (A) Overview of cerebellar differentiation protocol. Representative images of key stages are shown above (from left to right); undifferentiated iPSCs, Day 21 EBs, Day 21 EB cryosections immunostained for GBX2 (green) and EN1 (red), with nuclear Hoechst staining (blue), Day 35 organoid cryosections immunostained for KIRREL2 (red), with nuclear Hoechst staining (blue), and Day 60 organoids immunostained for Calbindin (green), TUJ1 (red), with nuclear Hoechst staining (blue). Scale bars are 1000 μM (first two panels) and 150 μM (all other panels). (B) mRNA levels displayed as fold-change values relative to undifferentiated iPSCs. Samples were calibrated relative to *ACTB*. *EN1* (p-value 0.0346), *GBX2* (p-value 0.1147), *EN2* (p-value 0.0087) and *OTX2* (p-value 0.6429). Results are shown from 5-6 independent differentiation experiments, Mann-Whitney test. (C) Quantification of EN1 and GBX2 immunostaining in day 21 EB cryosections as assessed by percentage of immunoreactive nuclei divided by total nuclei. Graphs show mean with SD from 4-7 independent differentiation experiments. (D) Day 35 mRNA levels of *OLIG2* (p-value 0.0023), *PTF1A* (p-value 0.0072), *ATOH1* (p-value 0.0004), and *KIRREL2* (p-value 0.4211). Violin plots show median and interquartile range. Data is shown from 8-9 independent differentiation experiments tested for significance using unpaired Mann-Whitney test. (E) The extent of polarized neuroepithelia was assessed counting rosettes per μM^2^ in cryosectioned organoids at day 35. Statistical significance was tested using a one-way ANOVA, with Sidak’s multiple comparison test. Untreated vs D7 p-value 0.9358, untreated vs D14 p-value 0.8587, untreated vs D21 p-value 0.0006. (F) Growth rate of organoids was significant over the course of differentiation (p-value 0.0001) and significantly increased by the addition of Matrigel (MG) from day 21onwards (p-value 0.0101) as assessed by two-way ANOVA, (Dunnet multiple comparison test). (G) Matrigel encapsulation resulted in a significant increase in *OLIG2* mRNA (p-value 0.0044), but non-significant changes in *PTF1A* (p-value 0.0857), *ATOH1* (p-value 0.2527) and *KIRREL2* (p-value 0.4893). Results are shown from three differentiation experiments, unpaired Mann-Whitney test.

### Encapsulation with Matrigel transforms growth of organoids

Matrigel (MG) has previously been used to simulate basement-membrane signalling and improve the growth of cortical organoids (Lancaster et al., 2017). To evaluate the effect of MG on the growth of cerebellar organoids and to determine the optimal potential responsiveness window of EBs to MG treatment, we encapsulated these at three different timepoints during differentiation. A significant difference (p-value 0.0006) in the number of rosettes indicative of polarized neuroepithelium was observed in EBs encapsulated on day 21 of differentiation in comparison to day 7 or 14 (ns, p-values 0.9358/0.8587) (Figure 1E). Following embedding on day 21, the average diameter of embedded organoids at day 35 was 1047.37 ± 367.47 μm in contrast to 752.71 ± 168.73 μm in unencapsulated organoids, indicating that the addition of MG to the culture setting resulted in divergence in growth (p-value 0.0101) (Figure 1F). In addition, excessive neuronal outgrowth was observed at the periphery of the organoids (data not shown). The addition of MG resulted in a significant increase in *OLIG2* mRNA expression (p-value 0.0044); however, it did not yield a significant difference in the expression of *PTF1A* (p-value 0.0857), *ATOH1* (p-value 0.2527) or *KIRREL2* (p-value 0.4893) (Figure 1G). Extensive and uniform expression of KIRREL2 was observed in both control and embedded aggregates on day 35 of differentiation, indicating no obvious change to VZ lineage-commitment upon MG treatment (Supplementary Figure S1).

### Single cell profiling of individual organoids reveals emergent cerebellar cell types and population heterogeneity

Organoids were allowed to mature for an additional 55 days at the air-liquid interface on transwell membranes before transcriptional profiling using scRNA-seq. Six organoids, each labelled with unique oligonucleotide hashtags, were sequenced as two hash-pools (2526 total cells prior to QC). Each hash-pool consisted of three control and three MG-embedded organoids, harvested on day 90 of culture. Clustering of raw and QC-filtered cells did not yield appreciable differences in population identity (data not shown). Given recent findings regarding the disparate and rapidly changing cell type-specific metabolic activity across developing neurons (Wizeman et al., 2019), conventional mitochondrial gene content QC cut-offs were discarded in favour of including all singlets regressed for mitochondrial content. Mitochondrial gene expression, unique molecular identifiers (UMIs) and gene counts are shown in Supplementary Figure S2, indicating near uniform mitochondrial and gene content across samples and hash pools, respectively.

Robust representation of equivalent populations was evident in both treatment groups following integration (Figure 2A). Optimal clustering to distinguish between cerebellar cell types was determined by visualizing separation of related clusters in pooled data at increasing resolutions using the ClusTree package (Supplementary Figure S3) and manual curation of the cluster biomarkers with canonical murine cerebellar markers. Following integration of treatment cohorts to identify common sources of biological variation, UMAP projection of canonical correlates at a resolution of 0.8 distinguished 12 distinct populations mutually occupying common transcriptional space (Figure 2B, C).

**Figure 2.**
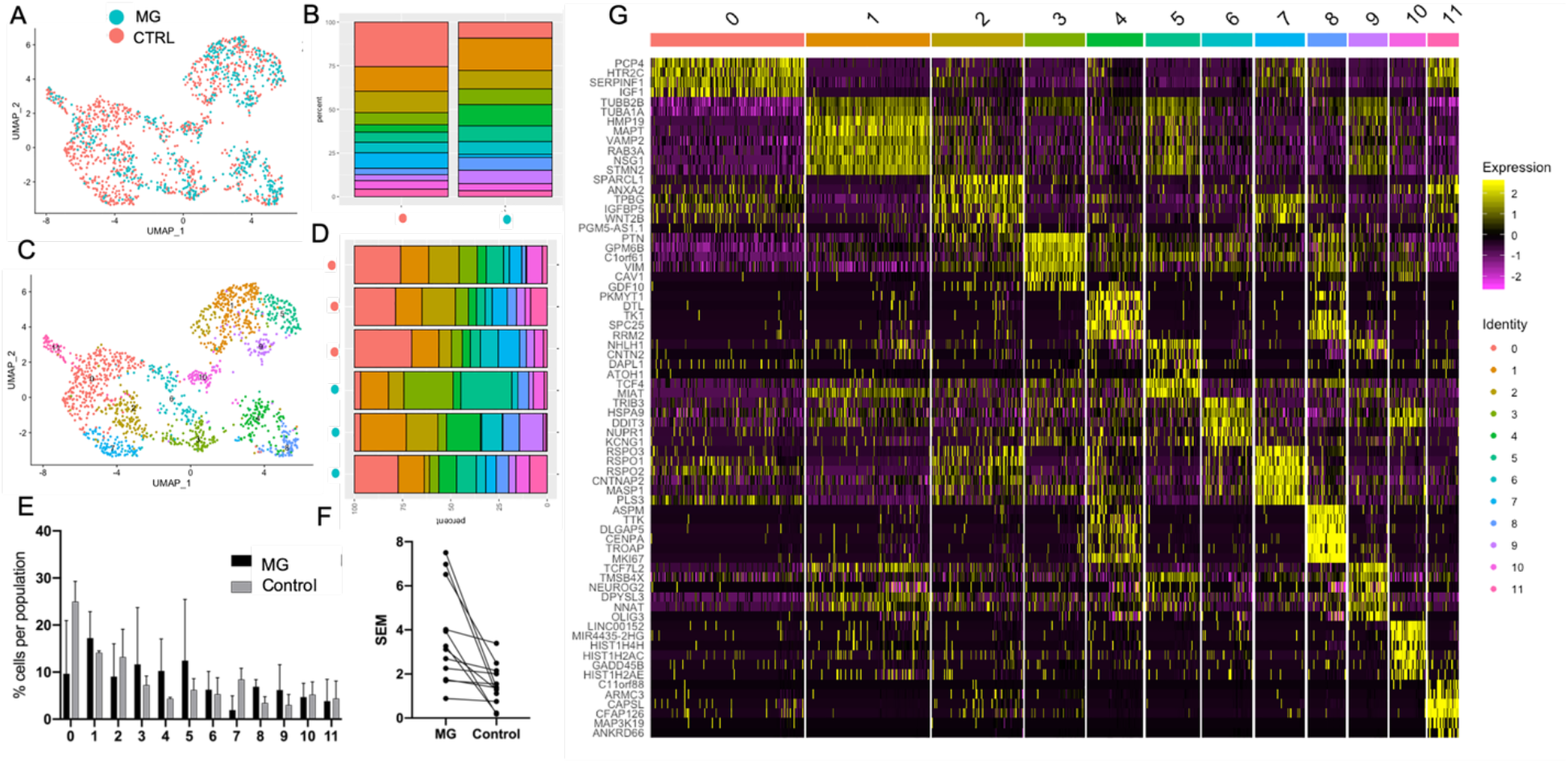
Characterisation of cerebellar organoids reveals population heterogeneity. (A) UMAP plot after canonical correlation analysis (CCA) illustrates representation of cells across treatment conditions (pink – control, teal – MG treatment from day 21). (B) Visual depiction of organoid heterogeneity, colored by population, shows representation of cell types across treatment groups. Left (pink): control (CTRL), right (teal): MG-embedded. (C) UMAP plot of all cells showing 12 populations following separation after CCA at resolution 0.8. (D) Visual depiction of organoid heterogeneity, colored by population, shows representation of cell types across individually-barcoded organoids. Top three samples (teal): MG-embedded, bottom three samples (pink): control. (E) Population composition shown by percentage, error bars depict mean/SEM. Population 0 comprised 9.67 ± 6.52 vs 24.98 ± 2.49 % of the total organoid composition, in encapsulated vs control, respectively. Population 1 (17.21 ± 5.65/14.12 ± 0.23%), Population 2 (9.07 ± 6.96/13.23 ± 3.39%), Population 3 (11.64 ± 12.09/7.25 ± 1.12 %), Population 4 (10.24 ± 6.82/4.34 ± 0.18 %), Population 5 (12.45 ± 12.99/6.22 ± 1.39%), Population 6 (6.24 ± 3.91/5.32 ± 2.01 %), Population 7 (1.92 ± 3.03/8.44 ± 1.38%), Population 8 (6.89 ± 1.54/3.46 ± 0.76%), Population 9 (6.18 ± 5.40/3.01 ± 1.30%), Population 10 (4.67 ± 2.94/5.23 ± 1.55%) and Population 11 (3.83 ± 4.67/4.40 ± 2.15). (F) Variability across control groups is shown by expressly visualizing SEM per population. (G) Heatmap depicting the top six differentially expressed genes per cell population, filtered by log fold-change values.

The majority of populations were represented across each organoid and treatment condition (Figure 2B, D). However, population heterogeneity per organoid was greater in samples embedded in MG compared to controls (Figure 2D-F). We employed differential expression testing to identify the top common genes driving separation of common populations across the integrated cohorts (Figure 2G, Supplementary Tables S1-12, and Supplementary Figure S4A, B). Population 0 (Figure 2 H, pink; top genes *PCP4, HTR2C, SERPINF1, IGF1, TUBB2B, TUBA1A*) expressed markers consistent with fourth ventricle derivatives and the hindbrain CP including *TTR, ID1, SPARC* and *BMP7*, which are known to be actively secreted from this region (Grimmer and Weiss, 2008). Population 1 (orange; top genes *HMP19, MAPT, VAMP2, RAB3A, NSG1 & STMN2)* expressed *TCF7L2*, recently identified to mark the RL-derived GCs and glutamatergic DCN (Gupta et al., 2018). Population 1 also expressed *LHX9*, a marker of ventral RL-DCN, and *DCX*, which is expressed as DCN neurons undertake migration to the nuclear transitory zone (Rahimi-Balaei et al., 2018). Population 2 (beige; top genes *SPARCL1, ANXA2, TPBG, IGFBP5, WNT2B, PGM5-AS1.1*) expressed astroglial and RP markers including *KRT18, IGFBP5, SPARCL1* and *PLTP*. Population 3 (lime; top genes *PTN, GPM6B, C1orf61, VIM, CAV1, GDF10*) exhibited expression of the characteristic Bergmann glia marker *FABP7*. Interestingly, we noted expression within this population of *PTPRZ1*, recently shown to mark Bergmann glia (Carter et al., 2018), and also *S100β*. Population 4 (green; top genes *PKMYT1, DYL, TK1, SPC23, RRM2, NHLH1*) also expressed *PCNA, MKI67* and *TOP2A* consistent with proliferative GCs. Population 5 (mint; top genes *NHLH1, CNTN2, DAPL1, ATOH1, TCF4, MIAT*) additionally expressed GC precursor markers *NEUROD2, NEUROD6, PAX3, MEIS1, DCC* and *NFIB* consistent with RL derivatives. Population 6 (teal; top genes *TRIB3, HSPA9, DDIT3, NUPR1, EIF1 & KCNG1*) also expressed *DLK1* and *CXCL14* consistent with PCs, which secrete CXCL14 to guide GC migration, during development (Park et al., 2012). Population 7 (blue; top genes *RSPO3, RSPO1, RSPO2, CNTNAP2, MASP1, PLS3*) also strongly expressed the genes *SPARC, S100B, LMX1A, HES1* and *TTR*, consistent with RP precursor cells. Population 8 (mauve; top genes *ASPM, TTK, DLGAP5, CENPA, TROAP, MKI67*) exhibited markers consistent with proliferative GCs in G2M phase, including *ASPM* which sustains postnatal cerebellar neurogenesis (Williams et al., 2015). Population 9 (purple; top genes *TCF7L2, TMSB4X, NEUROG2, DPYSL3, NNAT, OLIG3*) also expressed *NEUROD1* and *NHLH1*, consistent with RL precursors. Population 10 (pink; top genes *LINC00152, MIR4435-2HG, HIST1H4H, HIST1H2AC, GADD45B, HIST1H2AE*) identity could not be determined, although expression of *GADD45B* and close proximity to other VZ clusters might suggest GABAergic interneurons. Population 11 (salmon; top genes *C11orf88, ARMC3, CAPSL, CFAP126, MAP3K19, ANKRD66*) also expressed cilia-related genes *DYNC1I2, DYNLRB2, IFT22, TMEM67, FUZ* (Leightner et al., 2013; Stauber et al., 2017). Markers for all prospectively-identified cell types were verified using *in situ* hybridisation data from the Allen Brain Atlas (ABA) (Supplementary Figure S4B).

Together, our initial analysis identified the majority of the major cerebellar neuronal cell types including RL precursors, GCPs, GCs, PCs, Bergmann glia, CP, RP and glutamatergic DCN. As expected, the strongest patterns of cell-cycle progression emerged in clusters putatively ascribed in identity as GCs, known to be the most rapidly proliferating cell types in the cerebellum. Predicted cell cycle dynamics are shown by UMAP coloured according to cell identity (Supplementary Figure S5A) or predicted cell-cycle stage (Supplementary Figure S5B). Canonical genes driving S-phase/G2M (Tirosh et al., 2016) are projected per population as a heatmap (Supplementary Figure S5C). Supplementary Figure 5D shows ridge plots with predicted cell-cycle scores. Given the well-defined biology regarding cell-cycle progression in the developing cerebellum (Wang and Liu, 2019), we did not regress out cell-cycle effects. Broadly, VZ-derivative markers were observed in populations 0, 2, 3, 6, 7 and 11, with RL-derivative markers highly evident in populations 1, 4, 5, 8 and 9 (data not shown).

### An unbiased classification strategy confirms assignation of cerebellar cell types across species

Following putative identification of cluster identities, we sought to validate our findings using independent, unbiased approaches. A meta-comparison with reported markers from the literature provided support for identification of cluster identity (Supplementary Figure S6), revealing enrichment for canonical marker genes (full gene lists for corresponding cell types in Supplementary Tables S1-S12).

Recently published 10x scRNA-seq data obtained from embryonic day E10 to postnatal day P10 murine cerebella represents the most current and comprehensive atlas of cerebellar cell types from early embryonic to late postnatal murine development, identifying 15 distinct cell types and comprising 48 clusters across ten developmental timepoints (Carter et al., 2018). We utilized a strategy to identify pairwise correspondences across human and murine data and project equivalent cell types across species (Figure 3A, B). Following 1:1 human to mouse orthologue conversion and anchor integration, UMAP projection showed clustering of human organoid and murine cerebellar cell types (Figure 3B), confirming a high degree of confidence in cluster and cell type assignation (Figure 3A). For example, both putatively-annotated human GC groups clustered mostly with murine GCs and the designated glutamatergic DCN group clustered alongside the murine glutamatergic DCN cluster. Of note, both human RL and GC precursor groups clustered most closely to the mouse DCN group, reflecting their close developmental origin. The ciliated groups from both studies clustered strongly together. The human clusters of PCs, vascularized RP, CP, ‘Cluster 10’ and RP groups (all VZ derivatives) clustered together amidst the murine astrocyte, glia and progenitor cell clusters. This is consistent with the idea that the RP is comprised of precursors and astroglial cells, and that these groups all share a close developmental origin. Human Bergmann glia clustered with the murine astrocyte cluster.

**Figure 3.**
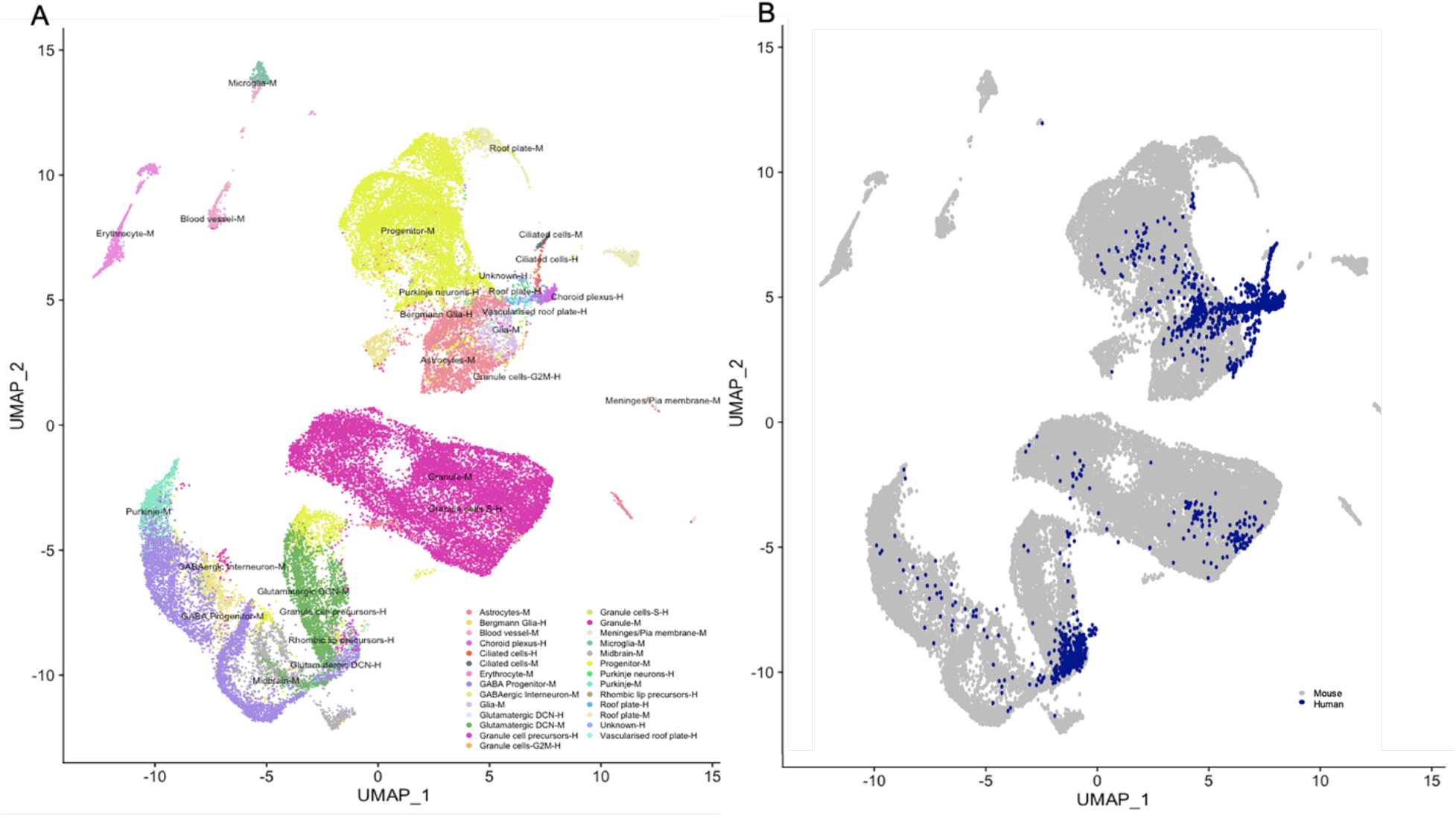
Human organoids recapitulate development of the cerebellum. (A) UMAP projection of integrated, human organoid data with murine developmental data, grouped by cell type identity reveals close proximity of human and murine cerebellar cell types. Human organoid-derived clusters are appended with −H. (B) Supporting UMAP displaying grouping by species.

To compare the maturity of human cell populations with specific developmental ages of the murine data, we projected temporal metadata onto the murine cell identity (Supplementary Figure S7). We observed overlap of the murine E15-P10 granule cells with the human S-phase GC cluster and E16-P10 murine cells with the human G2M-phase GC cluster (Supplementary Figure S8A-C). Murine glutamatergic DCN at E11-E17 clustered most strongly with the human glutamatergic DCN cluster (Supplementary Figure S8D, E). Human VZ derivatives overlapping with E13-E13 murine glia, E13-E17 astrocytes, and E10-E17 progenitor clusters respectively (Supplementary Figure S8 D,E). Thus, using the murine cerebellum as a close developmental blueprint, most signatures indicate a mixture of mid-late embryonic temporal maturity, suggesting that the cerebellar organoids recapitulate developmental stages of the normally developing cerebellum. An exception to this was overlap of human GCs with murine GCs of postnatal maturity, suggesting that this cell type was more mature than its counterparts.

We then employed a separate method to assess how effective the human to mouse cell type assignment was by utilizing the murine data as a reference dataset. Using this approach, we independently predicted murine cell type relatedness in human organoid clusters with a high degree of overlap (Supplementary Figure S9A). Supplementary Figure S9B shows canonical cell type marker expression in human organoid cells grouped by their predicted murine counterpart labels. Cluster annotation identified the murine counterparts present in the human cerebellar organoids with a high degree of concordance.

### Differential expression and trajectory reconstruction analysis reveals gross effect of Matrigel encapsulation on organoid differentiation

Following assignation of cell type clusters, we performed pseudo-time reconstruction to examine whether we could identify a developmental progression of maturing cell types. Trajectory reconstruction in pooled data showed a pattern reminiscent of the developmental cellular phylogeny of the cerebellum; cellular trajectory colour-coded by cell-type shows progression from primitive CP/RP cell types to RL/VZ precursors and subsequently to committed neuronal progeny bifurcating along the major glutamatergic/GABAergic lineages (Figure 4A). Major drivers of branching included a number of recently-identified markers involved with cerebellar specification (Haldipur et al., 2019; Wizeman et al., 2019) (Figure 4B). Branching topology was consistent following lineage reconstruction in separate treatment cohorts (Figure 4C), which also showed some altered branch dynamics (Supplementary Figure S13). Interactive pseudo-time reconstructions are available at https://plotly.com/~SamN1985/3 and https://plotly.com/~SamN1985/1 (control and MG respectively).

**Figure 4.**
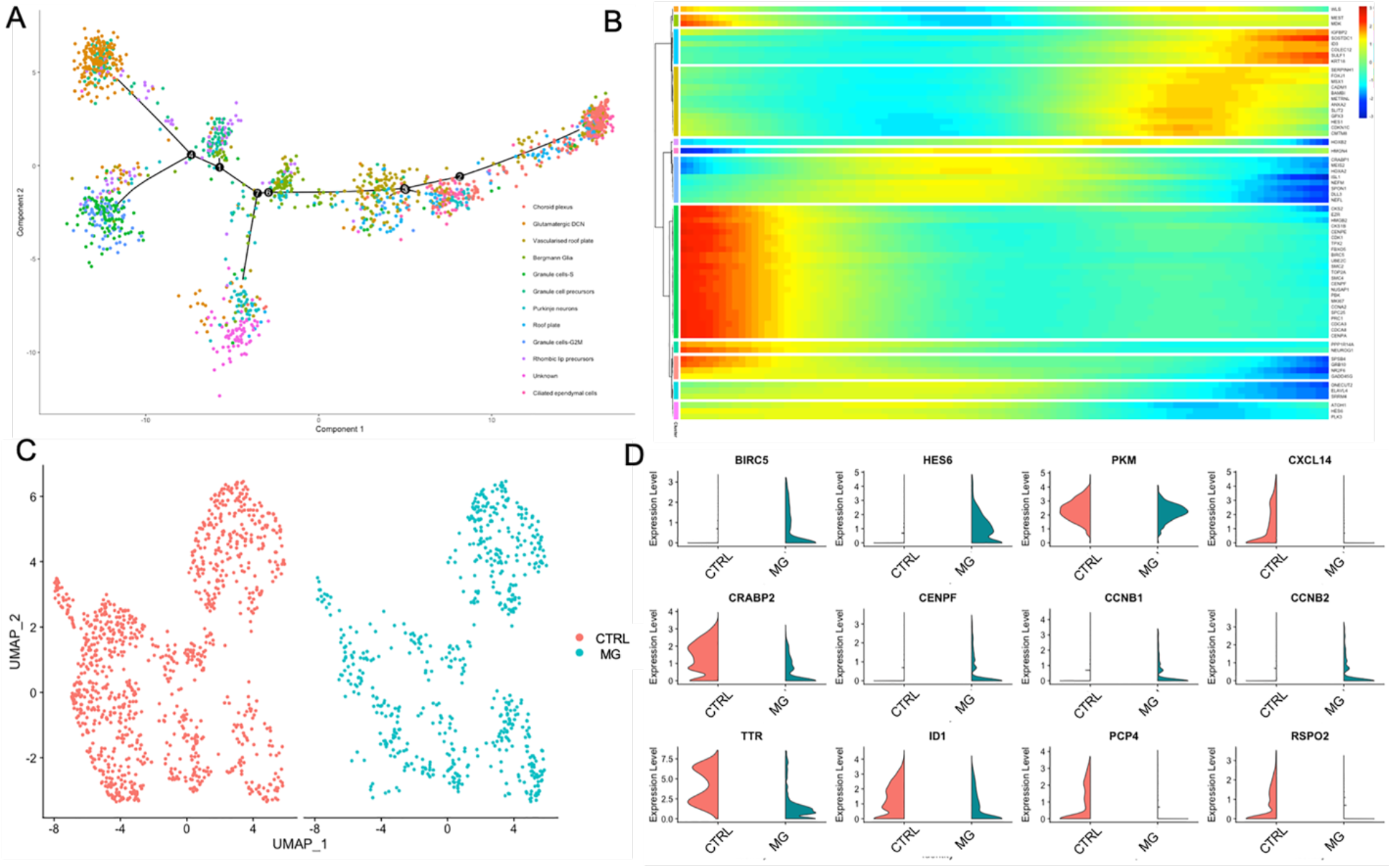
Organoid composition mimics development of the cerebellum, including basement-membrane contribution to RL derivatives. (A) Pseudo-time reconstruction of all hash-pooled organoids reveals a developmental trajectory akin to normal embryonic cerebellar development. (B) Pseudo-time plots show the contribution of canonical drivers of cerebellar specification per cell cluster. (C) UMAP projections of control (CTRL) and Matrigel (MG)-embedded organoids for differential testing. (D) Top differentially-regulated genes following MG encapsulation (p-value <0.05).

To explore embedding-related differences further, differential-expression testing between control and MG-embedded cohorts was performed. 46 genes were significantly differentially-regulated following MG encapsulation (36 genes FDR <0.05; 6 up, 30 down) as a cohort relative to control organoids (Supplementary Table S16). Gene-set overrepresentation analysis returned significantly altered Gene Ontology (GO) pathways *carboxylic acid biosynthetic process* (FDR 5.16E-05)*, organic acid biosynthetic process* (FDR 5.24E-05)*, regulation of angiogenesis* (FDR 6.21E-05)*, regulation of vasculature development* (FDR 1.1E-04)*, and tissue regeneration (*FDR 1.2E-04). Consistent with the transformation of growth and peripheral neuronal outgrowth following encapsulation, a number of genes concordant with survival, differentiation, and neurogenesis were among those significantly upregulated including *BIRC5*/Survivin, *HES6*, which promotes neurogenesis through the Notch pathway (Vilas-Boas and Henrique, 2010), *DCN*, which has been shown to regulate endothelial cell-matrix interactions during angiogenesis (Fiedler and Eble, 2009), and *PKM*, which inhibits proliferation during postnatal cerebellar neurogenesis (Tech et al., 2017) (Figure 4D). Furthermore, a number of hormones, chemokines and neuropeptides were amongst the genes differentially regulated with MG treatment. These include *TRH*, a hormone acting on receptors normally expressed in GCs and molecular layer interneurons (Watanave et al., 2018), *IGF1,* which is known to have pleiotropic effects in all neural cells including neurons, oligodendrocytes and astrocytes (Wrigley et al., 2017), *CXCL14*, a chemokine secreted by PCs which directs GC migration (Park et al., 2012), and *CRABP2*, which regulates RA production (Mosquera et al., 2018). In addition, several cell cycle-related genes were amongst the list of those differentially expressed (*CENPF, CCNB1, CCNB2, BIRC5, DCN*), suggesting that MG-encapsulation transforms organoids through modulation of cell-cycle dependant pathways. Furthermore, cell type-specific markers *TTR, ID1, DCN, PCP4, CXCL14, RSPO2* and *HES6* were differentially regulated following encapsulation (Figure 4D). *ID1* (downregulated in MG-embedded organoids, p-value 0.04755947) is transcriptionally active in embryonic ventricular and subependymal zones as well as being involved with postnatal development (Duncan et al., 1997). *ID* family members are known to regulate differentiation and their presence supports the detection of heterogeneity across treatment conditions. *TTR* and *PCP4* distinguish VZ precursors, while *DCN* is normally restricted to RL-derivatives.

Following several independent lines of cluster identity confirmation (Figure 3) and marker enrichment (Figure 4D), we observed a consistent bias in population composition in the MG-encapsulated cohort, towards cell-type identities arising from the RL (Figure 2B,E and F). To more clearly illustrate the relative contribution of RL/VZ derivatives, we performed retrospective re-clustering at a resolution of 0.09, distinguishing clusters comprising RL/EGL/VZ populations (Supplementary Figure S10A). Control organoids comprised 68.99±0.29% VZ derivatives compared with MG-embedded organoids, which showed 46.97±10.43% VZ derivatives (Supplementary Figure S10D), indicating a considerable bias away from the VZ following embedding. Chi-square testing revealed consistent VZ/RL bias following treatment at both clustering resolutions (Supplementary Figures S11 and S12). Validation of RL/EGL/VZ clustering is shown by enrichment of genes driving separation at this resolution (Supplementary Figure S10B), top markers (Supplementary Figure S10C) and genes distinguishing populations (Supplementary Tables 13-15).

### Population-level differential expression testing reveals cell-type specific responses to Matrigel

To evaluate the effect that MG encapsulation had on specific cerebellar populations (Figure 5A), gene over-representation tests were performed on differentially expressed genes (Figure 5B-C, FDR <0.05) across treatment conditions in equivalent populations. A list of differentially regulated genes is provided for each population comparison in Supplementary Tables S17-S28. GO-enriched pathways (Figure 5D) were examined for each population following treatment. Population 0 (CP) following MG encapsulation exhibited significantly altered regulation of *‘Steroid biosynthetic process’* (FDR 7.44E-05) and *‘Cholesterol biosynthetic process*’ (FDR 7.44E-05). This is interesting as cholesterol release from epithelial cells of the CP epithelia has been described to play an important role in cholesterol homeostasis in the cerebrospinal fluid (Kant et al., 2018). Population 3 (Bergmann glia) showed significant changes in ‘*Retinoic acid metabolic process’ (*FDR 1.68E-04). Notably, release of retinoic acid by radial glial cells has shown to be instrumental for numerous developmental processes including formation of the blood-brain barrier (Mizee et al., 2013). Population 4 (GC precursors) showed differential regulation of ‘*DNA packaging’* (FDR 9.27E-08), ‘*nucleosome assembly’* (FDR 2.00E-06), *‘chromatin remodelling at centromere’* (FDR 1.25E-05) and ‘*telomere maintenance via semi-conservative replication’* (FDR 1.78E-04). Population 6 (PCs) displayed enrichment for pathways ‘*Intrinsic apoptotic signalling pathway in response to endoplasmic reticulum stress’* (FDR 5.68E-09), *‘response to starvation’* (FDR 5.03E-05) and *‘monocarboxylic acid biosynthetic process’* (p-value 1.71E-05). PCs express high levels of monocarboxylate transporter 2, an integral component of the lactate transport system required for energy metabolism and neuronal activity (Bergersen et al., 2001). Overall, the relative effect of MG encapsulation resulted in distinct responses in the various cerebellar populations. This was most notable in RL-derivatives, where encapsulation seemed to influence cell-cycle dynamics in GCs. MG encapsulation altered a number of cell type-specific pathways known to be operational during normal development, however, the most noteworthy effect on the organoids was the amplification of RL-derivatives driving expansion of the GC population. This is especially interesting given the fact that GCs were proportionally under-represented compared to the physiological composition of the cerebellum.

**Figure 5.**
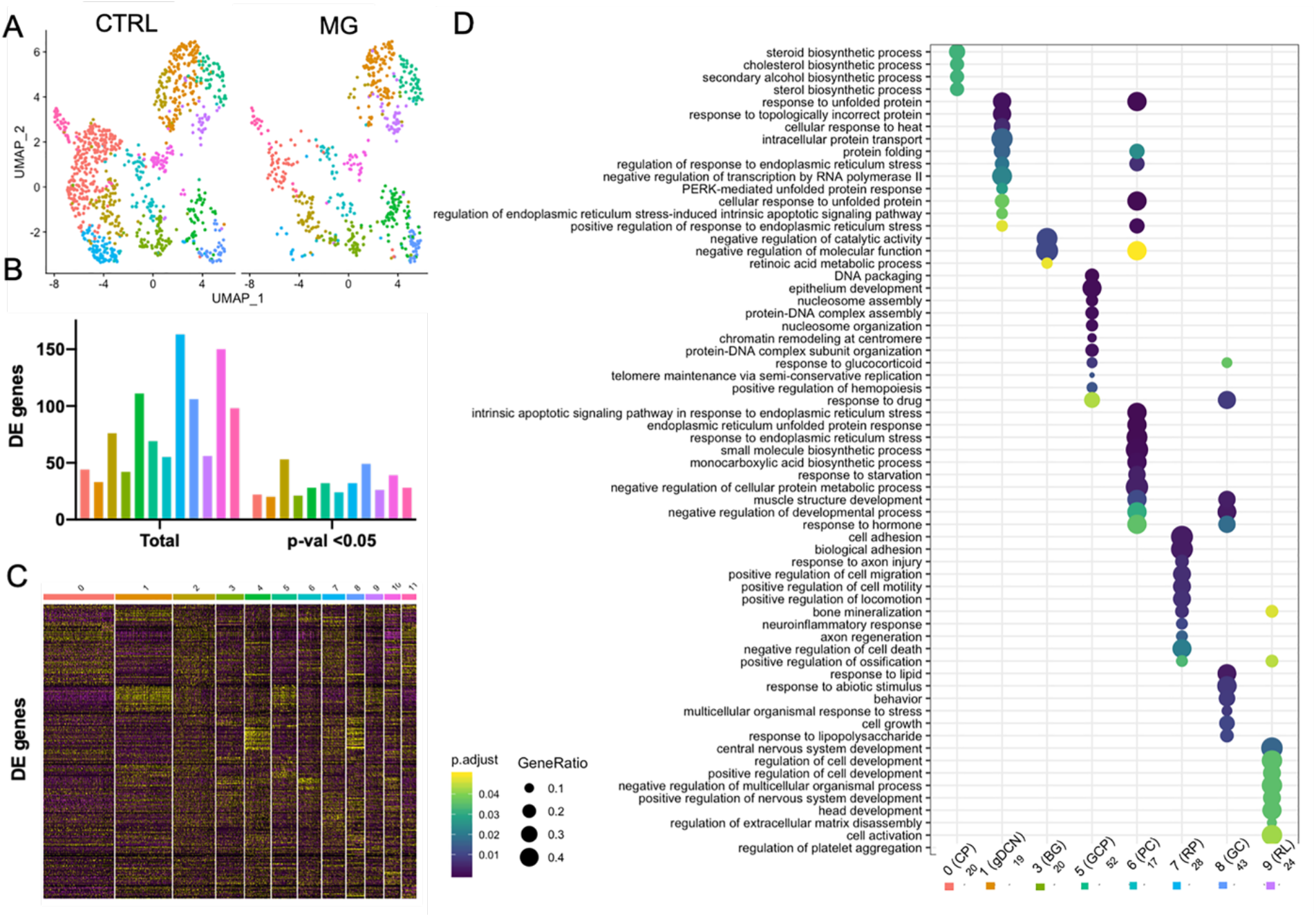
Cerebellar populations show distinct responses to Matrigel encapsulation. (A) UMAP projections outlining equivalent populations from control (CTRL) and Matrigel (MG)-embedded cell groups for differential-expression testing. (B) Graph depicting numbers of total (left) and significantly changing genes per cell population. P-value <0.05 accepted as significant. (C) Heatmap depicting all differentially-regulated genes between untreated and MG-embedded organoids. (D) Dot plots showing the top differentially expressed GO pathways per cell population. The number of genes considered for each population is listed on the x-axis below cell labels.

## DISCUSSION

We have developed a robust and reproducible protocol for the generation of three-dimensional cerebellar organoids from hiPSCs. By employing scRNA-seq we demonstrate that the cerebellar organoids contain transcriptionally-discrete populations encompassing all major cerebellar neuronal cell types including RLPs, GCPs, PCs, GCs, RP, CP, Bergmann glia, and glutamatergic DCN, and recapitulate properties of the normally developing cerebellum; this includes proximally-located territories in which adjacent signalling is required for cerebellar maturation and development. This model system has promising implications for the study of cerebellar development and disease most notably by obviating the necessity for murine co-culture, thus providing xeno-free conditions that represent a more biologically relevant culture setting, which may also be more therapeutically tractable.

Cell-autonomous processes, as well as the local microenvironment drive specification and development of the cerebellum. A popular method to promote sustained growth of cortical organoids has been the addition of MG to the culture setting, promoting differentiation through exposure to molecules present at the basement membrane (Lancaster et al., 2018). In the cerebellum the ECM-enriched basement membrane is directly below the pial meningeal layer, which provides an anchor point for endfeet of radial processes from VZ precursors and also forms a barrier to migrating neurons (Halfter and Yip, 2014; Nguyen et al., 2013). In both cortical and cerebellar development, these processes provide a scaffold that facilitates neuronal migration and correct layer formation. An example of this was shown by (Takeuchi et al., 2015), demonstrating the necessity for type IV collagen which controls GC axon formation by regulating the integrity of the BM. In agreement with these observations, our data show that migratory and proliferative activity is significantly altered in MG-embedded cerebellar organoids. However, encapsulation also decreased the reproducibility of generating cerebellar cell types, with embedded organoids showing a greater degree of variability in cell-type composition. Interestingly, this heterogeneity comprised a bias towards RL-derivatives, including glutamatergic DCN and GCs. This is especially intriguing, given that previous cerebellar differentiation protocols have been reliant on co-culture with isolated murine RL progenitors or GCs to provide necessary factors for growth and maturation of the VZ component (Ishida et al., 2016; Muguruma et al., 2015; Sundberg et al., 2018). Our population-level differential expression analysis suggests that GCs were especially-sensitive to the addition of MG, showing a greater degree of responsiveness as marked by increased numbers of significantly-altered genes. This suggests that MG encapsulation of cerebellar-patterned organoids may provide a viable alternative to costly, time-restrictive rodent RL isolations that hamper downstream analyses of the differentiated human cells.

An exciting potential for organoid-based protocols is the fact that neighbouring regions can influence the development of adjacently-located tissues. For example, both RP epithelium and CP epithelium are required for full development of the RL *in vivo* (Broom et al., 2012; Chizhikov et al., 2006). It has been suggested that diffusible factors such as retinoic acid influence development of the cerebellum, following secretion from the CP (Krizhanovsky and Ben-Arie, 2006). Our organoids, which do not require murine feeder layers, contained CP, RP, ciliated cells, RL precursors, glutamatergic DCN, GCs, Bergmann glia and PCs. This provides strong evidence that organoids transit through stages akin to physiological embryonic cerebellar development. Furthermore, following MG encapsulation, organoids underwent significant changes in physiologically-relevant pathways including Bergmann glia displaying altered *retinoic acid biosynthetic processes* and CP cells *steroid & cholesterol biosynthetic processes*. Heavy overrepresentation of cell-cycle and DNA synthesis-related genes and pathways in RL-derivatives indicates that MG encapsulation drove expansion of this cell type *in vitro*.

The presence of CP cells within the organoids is exciting given the functional implication for this model. The CP is a non-neuronal structure, comprised of both blood vessels and epithelial cells, located adjacent to the RL. The CP forms one of the interfaces of the blood-brain barrier and is involved in the production of cerebrospinal fluid, as well as IGFII, GDNF, and TGFα (Krizhanovsky and Ben-Arie, 2006). One possibility regarding the heterogeneity between treatment conditions is that MG-embedding inhibits formation of the CP, which would, in turn, attenuate formation of the RL, mediated by BMP7, a mechanism similarly described *in vivo* (Krizhanovsky and Ben-Arie, 2006). While several other organoid protocols are introducing vasculature, the presence of CP with endothelia suggests that our organoids may possess an intrinsic capability to produce some degree of vascularisation.

Our integration with existing single-cell data from the developing mouse cerebellum suggests that the human organoids resembled mid-to-late murine embryonic development. Interestingly, while the majority of our annotated human cell types clustered with their murine counterparts, human PCs clustered more closely to murine progenitors and astroglia, suggesting that by day 90 organoid-derived PCs were still developmentally immature, compared with murine PCs. In further support of this, we did not detect appreciable levels of *SHH*. Findings in the literature suggest that PCs and GABAergic DCN arise from the VZ prior to dependence on *SHH* signalling (Huang et al., 2010), consistent with the presence of immature PCs in the organoids that are still undergoing maturation, potentially in suboptimal culture settings. Further comparison of the organoid-derived PCs to murine PC transcriptomes may yield some clues about more supportive trophic settings. Interestingly, the unknown population (cluster 10) clusters proximally to PCs, raising the interesting possibility that this represents a PC subtype or perhaps another GABAergic neuronal cell type.

hiPSC-derived organoid models offer unprecedented opportunities to model brain development and disorders and for therapeutic development. The cerebellar organoid model presented here will help to better understand the processes required for neuronal maturation in the current culture paradigms. Moreover, using patient-derived hiPSCs, this model is poised to provide novel insights into the pathogenic mechanisms underlying disorders of the cerebellum.

## MATERIALS AND METHODS

### hiPSC growth and maintenance

hiPSCs from a healthy 67-year-old female donor (AHO17-3) were used as previously described (Watson et al., 2018) between passage 25-35. Samples were tested and free of Mycoplasma using the MycoAlert kit (Lonza, Switzerland). hiPSCs were maintained on Matrigel-coated plates in a humidified CO_2_ incubator with mTeSR1 media changed daily (STEMCELL Technologies, UK). hiPSC were passaged upon reaching approximately 80% confluence using EDTA at 37°C for three to five minutes and plated in mTESR1 with 10 μM Y-27562 (Tocris, UK). Media without Y-27562 was supplemented the day following.

### Cerebellar organoid differentiation

Figure 1A provides an overview of the differentiation protocol. Briefly, embryoid bodies (EBs) were made by dissociating sub-confluent hiPSCs with TrypLE before allowing to aggregate in ultra-low attachment 96 well-plates (PrimeSurface MS-9096V, Sumitomo Bakelite, Japan) in growth factor-free chemically-defined medium in the presence of 7μg/ml Insulin (Sigma, USA), 50 μM Y-27562 (Tocris) and 10 μM SB431542 (Muguruma et al., 2015). Approximately 1.1E04 cells were deposited into each well. Two days later, media was supplemented with 50 ng/ml FGF2 (R&D Systems, USA). On day seven, one-third of the media was removed and supplemented with fresh media without growth factors. On day 14 full volume media replacement was performed. On day 21 of differentiation, individual EBs were embedded in undiluted Matrigel or transferred into differentiation media as free-floating aggregates (four EBs per well of a 24-well low-attachment plate). EBs were harvested on day 21 for cryosectioning/qPCR to validate acquisition of MHB fate. Medium was changed on day 28 and 35 before assessing expression of rhombic lip/ventricular zone markers. EBs were monitored for growth by photographing every two days and inspected for visual signs of differentiation including the presence of polarized neuroepithelia (rosettes). On day 35, individual EBs were transferred to transwell membranes (Millicell Cell Culture inserts, PTFE, 0.4 μM, Merck Millipore, USA) and allowed to further develop at the air-liquid interface. Organoids were harvested at multiple timepoints throughout for validation of successful cell-fate acquisition.

### Immunostaining

EBs and organoids were harvested for cryosectioning at various timepoints by fixing in 4% paraformaldehyde for 15 minutes at room temperature, followed by three washes of PBS. Samples were cryoprotected overnight in 20% sucrose/PBS before embedding in OCT and sectioning on a cryotome at 10 μM. Cryosections were blocked in 2% milk power/PBS for one hour at RT following permeabilization with 0.3% Triton-X100 for 10 minutes at RT. Sections were incubated with primary antibody diluted in blocking buffer ± 0.3% Triton-X100 overnight at 4°C followed by application of relevant fluorescent secondary-conjugated antibody (Alexa Fluor, Invitrogen). An identical preparation without the primary antibody was used as a negative control. Hoechst was applied for 5 minutes at RT to visualize nuclei. Immunostained sections were mounted in Prolong gold anti-fade (VECTASHIELD). A list of antibodies can be found in Supplementary Table S29.

### Quantitative real-time PCR

RNA was isolated from EBs/organoids on days 0, 21 and 35 using the RNeasy mini kit (Qiagen, Germany) according to manufacturer’s instructions including an on-column DNAse digestion step. cDNA was generated using the Superscript III First-strand synthesis kit (Invitrogen, USA). Quantitative real-time PCR was performed on an Applied Biosystems StepOne plus qPCR machine using Fast SYBR Green Mix according to the manufacturer’s specifications. No template and minus reverse transcriptase controls were used to test for specificity. Primers are listed in Supplementary Table S30. Relative fold-change of mRNA abundance was calculated using the ΔΔCq method. Samples were calibrated relative to *ACTB*. Results are shown from between 5 and 6 independent differentiation experiments. ΔCq values were tested for significance with P <0.05 accepted as significant, Mann-Whitney test.

### Imaging and quantification

Imaging was performed on mounted cryosections/whole-organoids using a Zeiss upright fluorescent microscope. CellProfiler was used to segment and identify immunopositive nuclei in fluorescently-labelled cryosection images. This was divided by the total number of nuclei as ascertained by visualisation with Hoechst. For rosette quantification, sample identity was blinded and polarized pockets were counted per section using an ImageJ cell counter plugin. Counts were normalized to total cryosection area covered by tissue. Organoid growth was assessed by measuring diameter using ImageJ and assessed for significance through two-way ANOVA. Formation of polarized neuroepithelial tissue was quantified in blinded samples and subject to statistical testing using one-way ANOVA.

### Statistical analysis

Statistical tests were performed with the PRISM software package (v8) or R. qPCR data is shown from multiple independent differentiation experiments, ΔCq values tested for significance using unpaired Mann-Whitney test. Rosette formation was assessed for statistical significance using a one-way ANOVA, with Sidak’s multiple comparison test. Growth rate of organoids was assessed by two-way ANOVA (Dunnet’s multiple comparison test). Chisquare testing was used to assess lineage-specific bias in SC data at both utilized clustering resolutions. P-value <0.05 accepted as significant for all statistical testing.

### Preparation of organoids for single cell sequencing

Following 90 days of differentiation, three organoids embedded in Matrigel (at day 21) and three unembedded organoids were manually dissected from transwell membranes and enzymatically digested with Accumax dissociation reagent (Sigma) with manual trituration. Organoids were resuspended in 0.01% BSA in PBS on ice and labelled with Hash-tagged Oligonucleotide (HTO)-conjugated antibodies for B2M/ATP1B3 at RT for one hour (HTO sequences and antibodies are listed in Supplementary Table 31). Samples were processed according to the standard 10x Chromium 3’ workflow and pooled before sequencing over two lanes (Illumina HiSeq 4000).

### Processing of single cell data

A total of 2,526 cells from two hashed pools were sequenced at an average of 53,751 reads per cell, with a median detection rate of 2,617 genes per cell. Hashed pools were demultiplexed using Seurat HTODemux with default options and 65% of the cells (1653) were classified as singlets. The SCTransform function in Seurat (v 3.2.1) was used to normalize UMI depth across cells using negative binomial regression followed by additional regression for mitochondrial proportion across all cells. Each hash pool was analysed separately and then also integrated and projected in UMAP space using canonical correlation analysis (CCA) and MNN (Mutual nearest neighbours). The ClusTree package and manual curation was used to visualize and determine optimal clustering parameters. Populations were identified through unsupervised clustering and projection of known cell-type specific markers using the FindConservedMarkers using default parameters (Love et al., 2014; McDavid et al., 2013; Trapnell et al., 2014). Organoid composition was projected by plotting the fraction of each population per organoid after integration/CCA + MNN projection. Expression of cluster-specific markers was validated through comparison to sagittal ISH sections of the developing murine brain (Allen Brain Atlas). Differential expression testing was performed using the FindConservedMarkers command using a Wilcoxon Rank Sum Test using identity classes comprising the two hash pools (Matrigel-embedded samples vs control). Mouse vs human comparisons were made by performing Integration and Label Transfer (Stuart et al., 2019) and incorporating high-confidence/1:1 human to mouse orthologues from Ensembl. Pseudo-time reconstruction was performed using Monocle. Code is available on Github repository SamN1985. A list or identities pertaining to cluster identity from murine dataset is listed in Supplementary Table 32. Data is available on GEO (GSE150153) and Stemformatics (<u>pending</u>).

### Pathway analysis

GO Term Biological process over-representation tests across all cell populations was performed using the *CompareCluster* function from the Bioconductor package ClusterProfiler (Yu et al., 2012). For each cell type population, differentially expressed genes between treatments with a p.value < 0.05 were included in the analysis and the background universe consisted of 17,125 genes that were expressed in a minimum of 3 cells with a UMI count >= 1.

## ACKNOWLEDGEMENTS

We thank M. Attar, H. Slawinski, and S. Revale for technical assistance and M. Cooper, L. Harris for their comments on the manuscript. S.N. is the recipient of a fellowship from the Oxford Nuffield Medical Trust. This work was supported by BrAshA-T, Action for A-T, the Rosetrees Trust, and the Wellcome Trust via core funding to the Wellcome Centre for Human Genetics (award 203141/Z/16/Z).

## AUTHOR CONTRIBUTIONS

S.N and E.B conceived the project. S.N performed differentiation experiments, qPCR, immunocytochemistry, sample preparation, scRNASEQ analysis and data visualisation. D.A assisted with scRNASEQ analysis, interpretation and data visualisation. F.C assisted with scRNASEQ analysis and interpretation. R.B assisted with scRNASEQ analysis and interpretation. S.N and E.B. wrote the manuscript with feedback from all authors.

## DECLARATION OF INTERESTS

The authors declare no competing interests.

## Notes

https://github.com/SamN1985/General-code

https://plotly.com/~SamN1985/3

https://plotly.com/~SamN1985/1

https://www.ncbi.nlm.nih.gov/geo/query/acc.cgi?acc=GSE150153

### Competing Interest Statement

The authors have declared no competing interest.

