## Supplementary figures for "Single-cell sequencing of human iPSC-derived cerebellar organoids shows recapitulation of cerebellar development"

Supplementary Figure S1

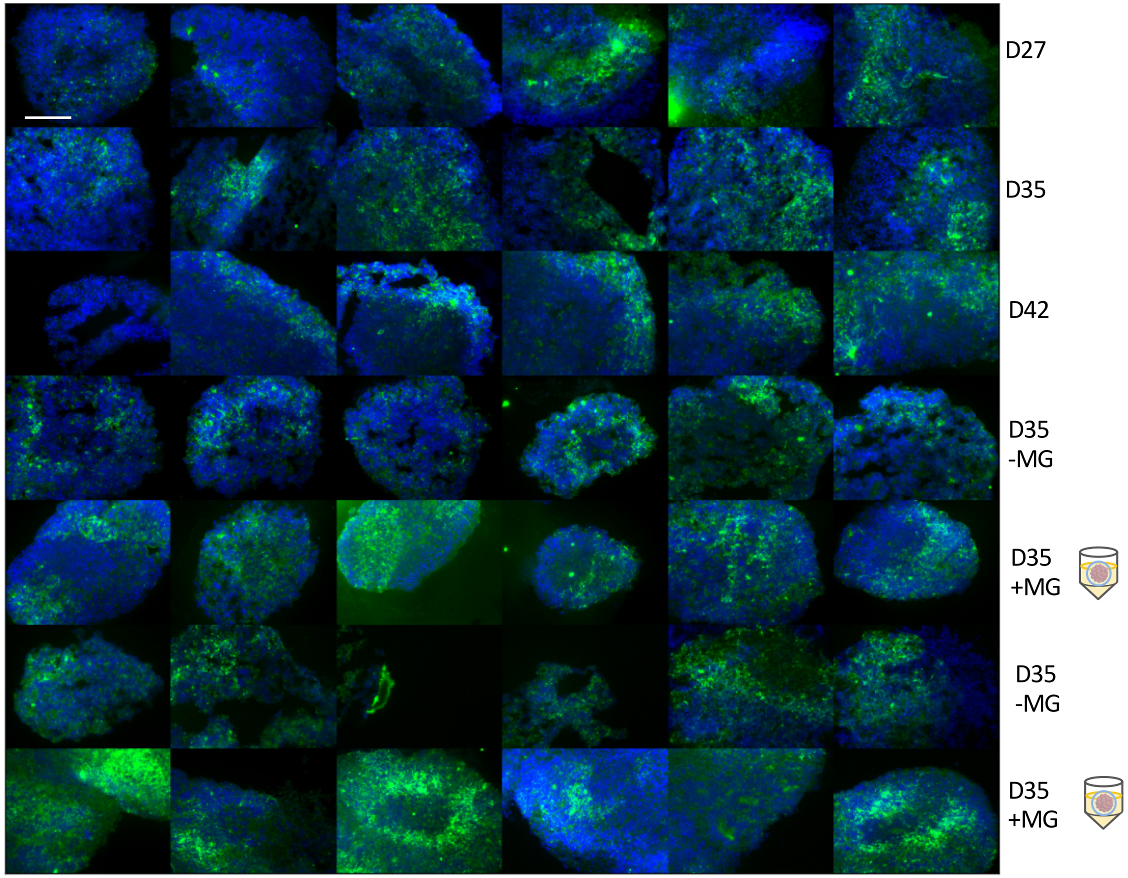


**Supplementary Figure S1 –** KIRREL2 (green) staining of organoids from three independent differentiation experiments. Top row, experiment 1 D27, Second row, experiment 1 D35. Third row, experiment 1 D42. Fourth row, experiment 2 D35. Fifth row, experiment 2, D35 with Matrigel embedding on D21. Sixth row, experiment 3, D35. Seventh row, experiment 3 D35 with Matrigel embedding on D21. Scale bar is 150 μm. Hoechst stains the nucleus. Related to Figure 1.

#### Supplementary Figure S2

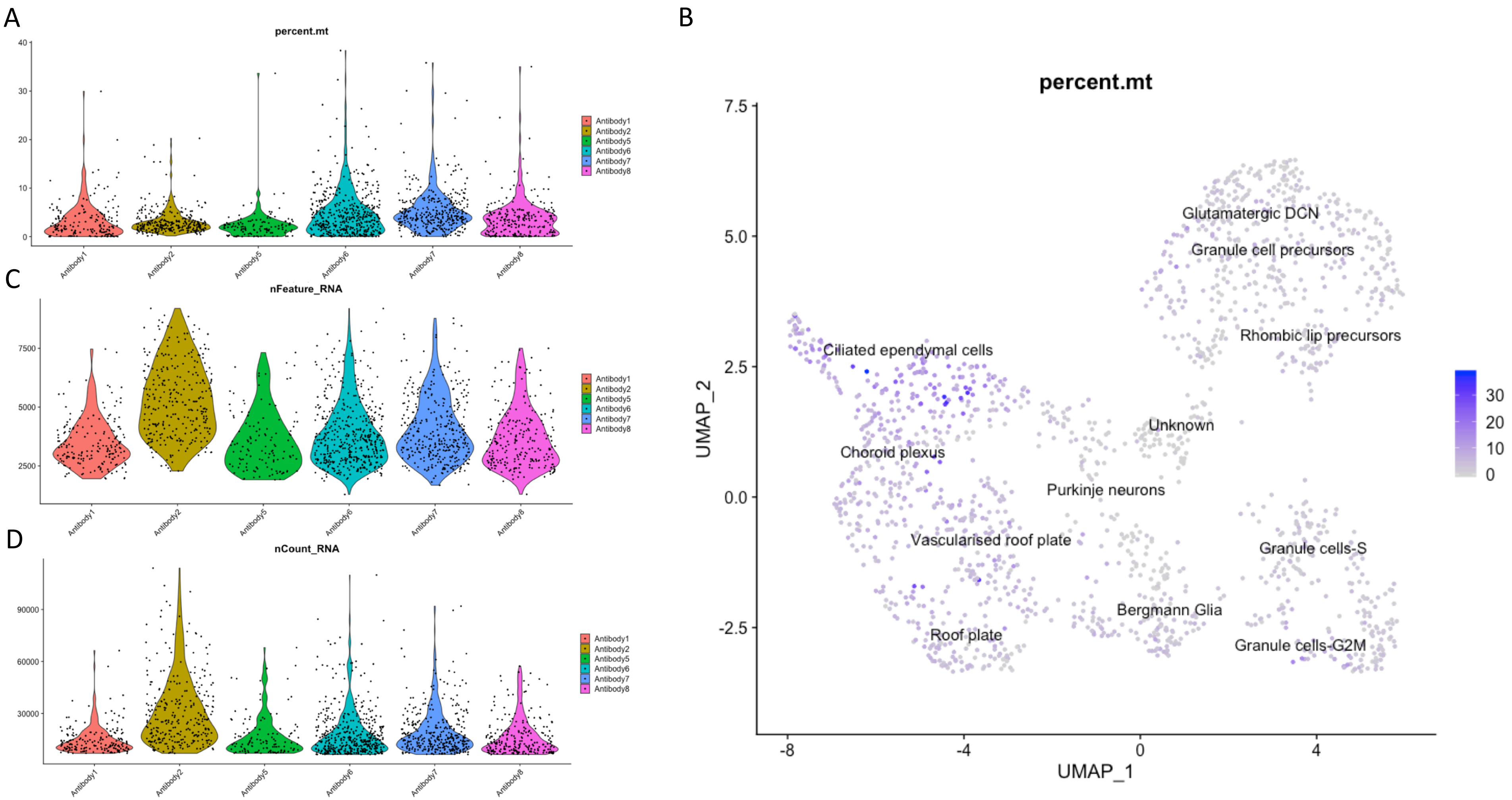


**Supplementary figure S2 –** Violin plots depicting (A) mitochondrial gene expression, (B) gene content and (C) UMI distribution in samples/hash pools. D) Mitochondrial content projected in UMAP space. Related to Figure 2.

Supplementary Figure S3

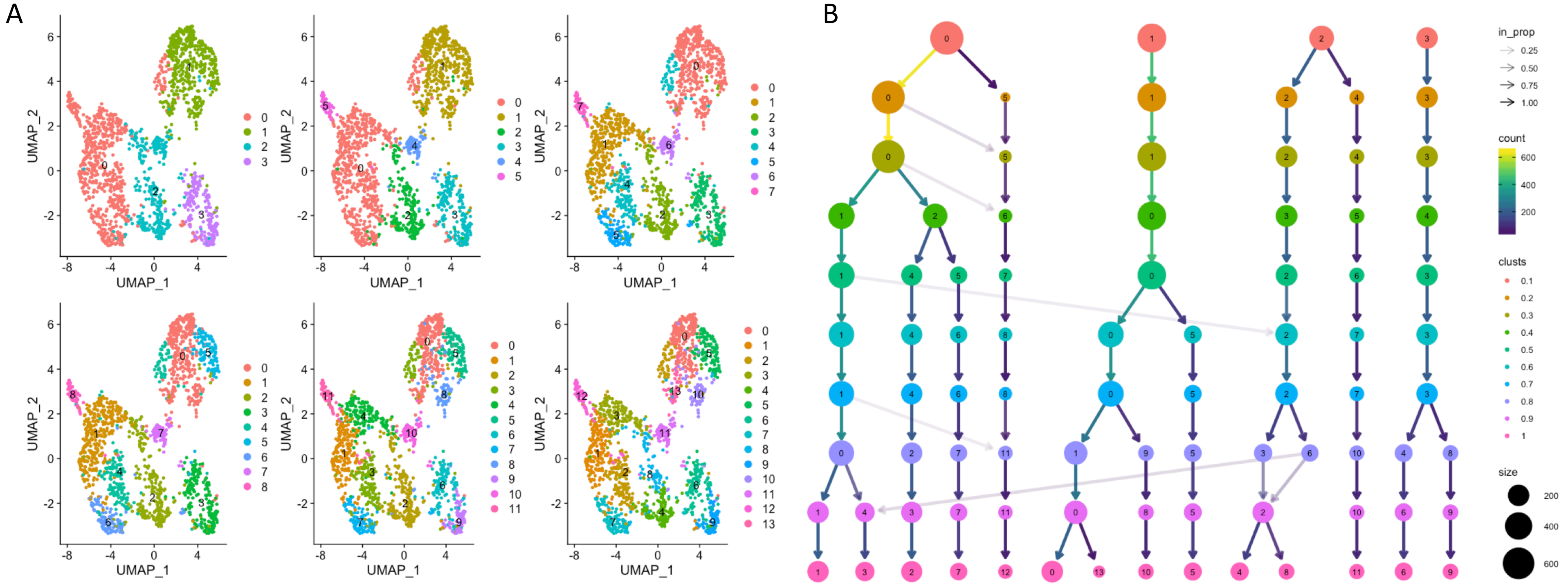

**Supplementary figure S3 –** A) UMAP and B) ClusTree depiction of cluster identity at increasing resolutions. Related to Figure 2.

Supplementary Figure S4

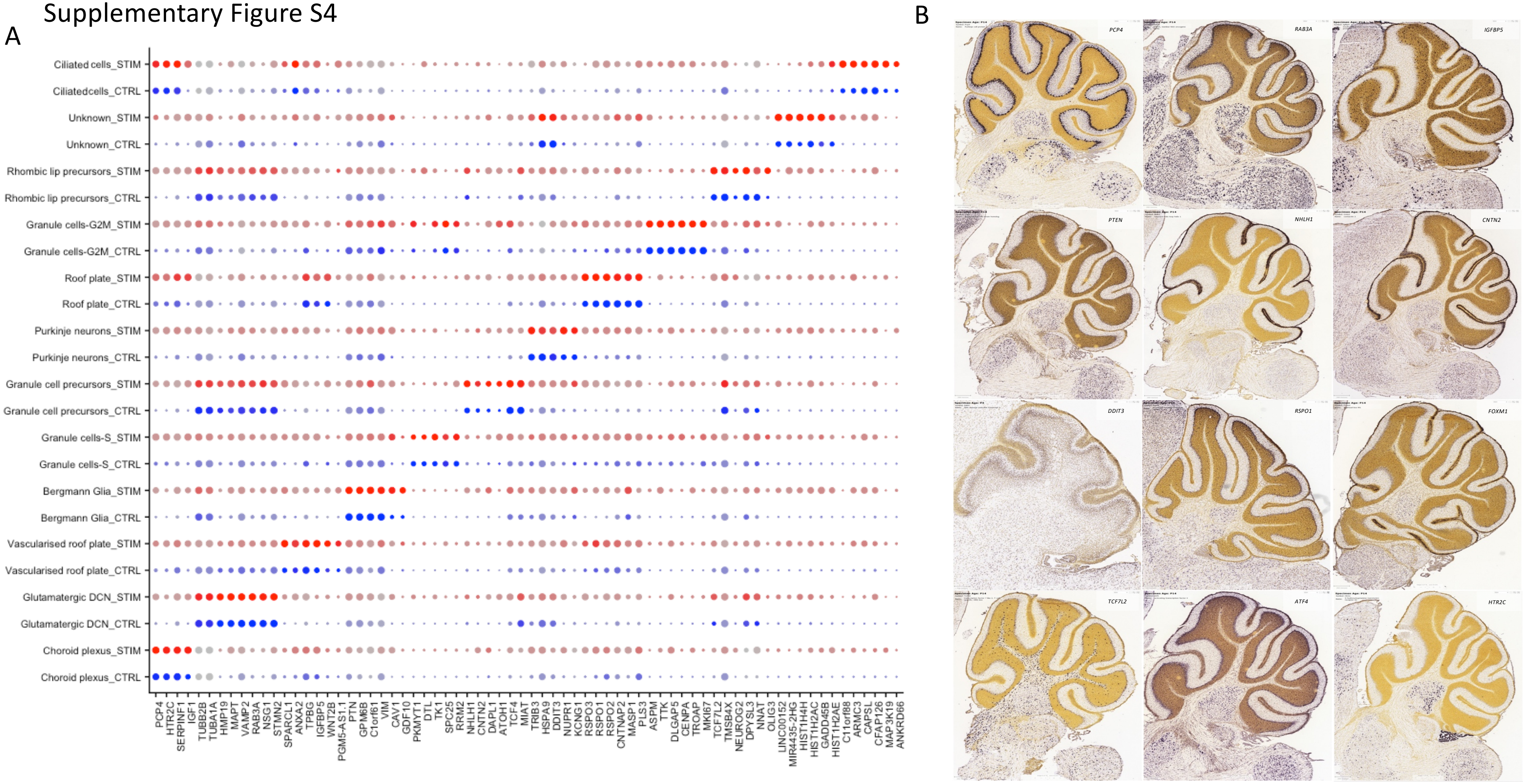


**Supplementary figure S4 –** A) Dot-plot showing expression of key markers in identified populations in Matrigel-embedded (STIM) and control samples (CTRL). B) Corresponding murine ISH sections from the ABA. Related to Figure 2.

### Supplementary Figure S5

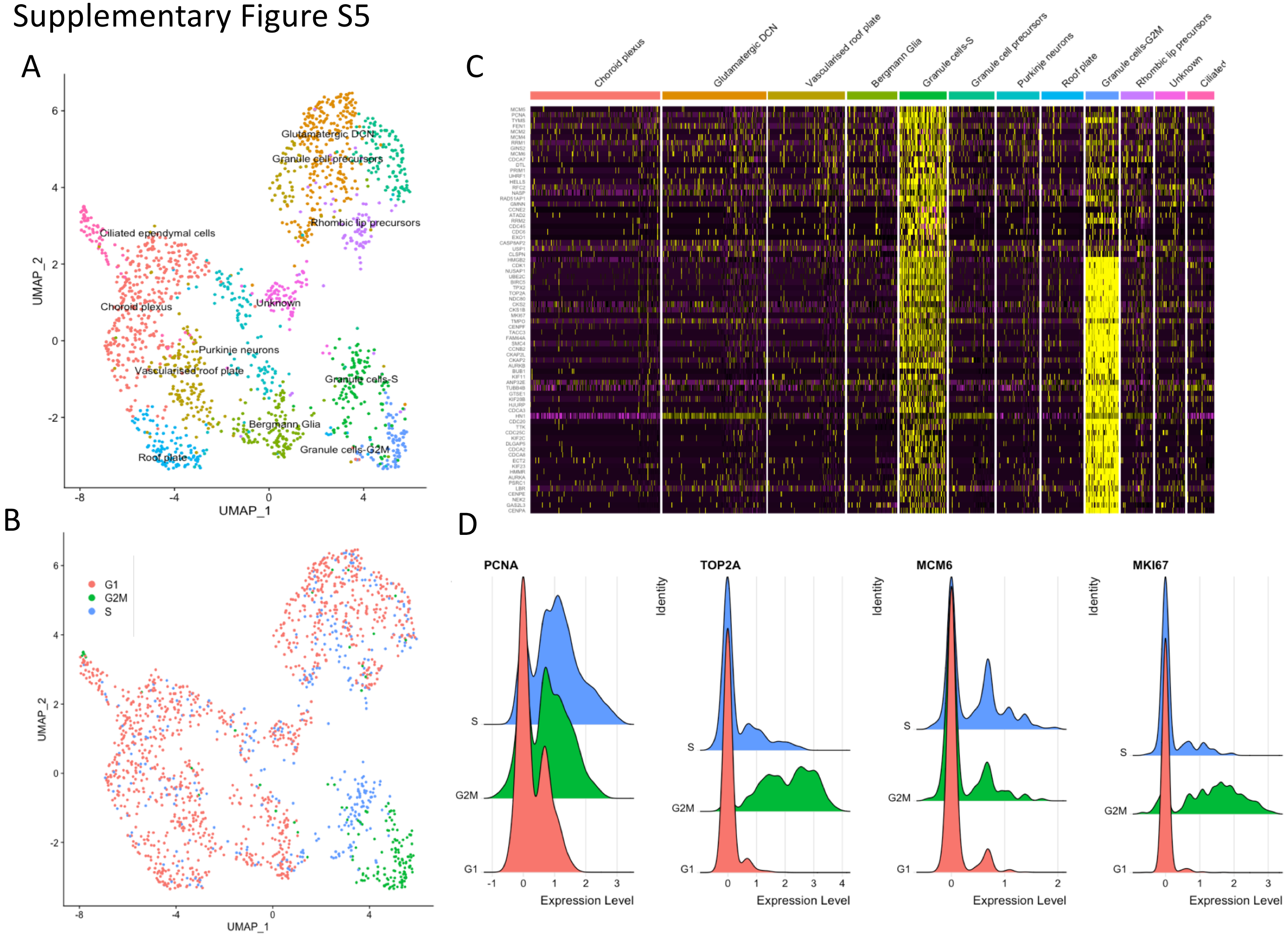


**Supplementary figure S5 –** A) UMAP projection of human organoids by cell type. B) UMAP projection of human organoids coloured by predicted cell-cycle phase. C) Heatmap of cell cycle-related genes. D) Ridge-plots depicting critical cell-cycle genes *PCNA, TOP2A, MCM6* and *MKI67* expression. Related to Figure 2.

Supplementary Figure S6

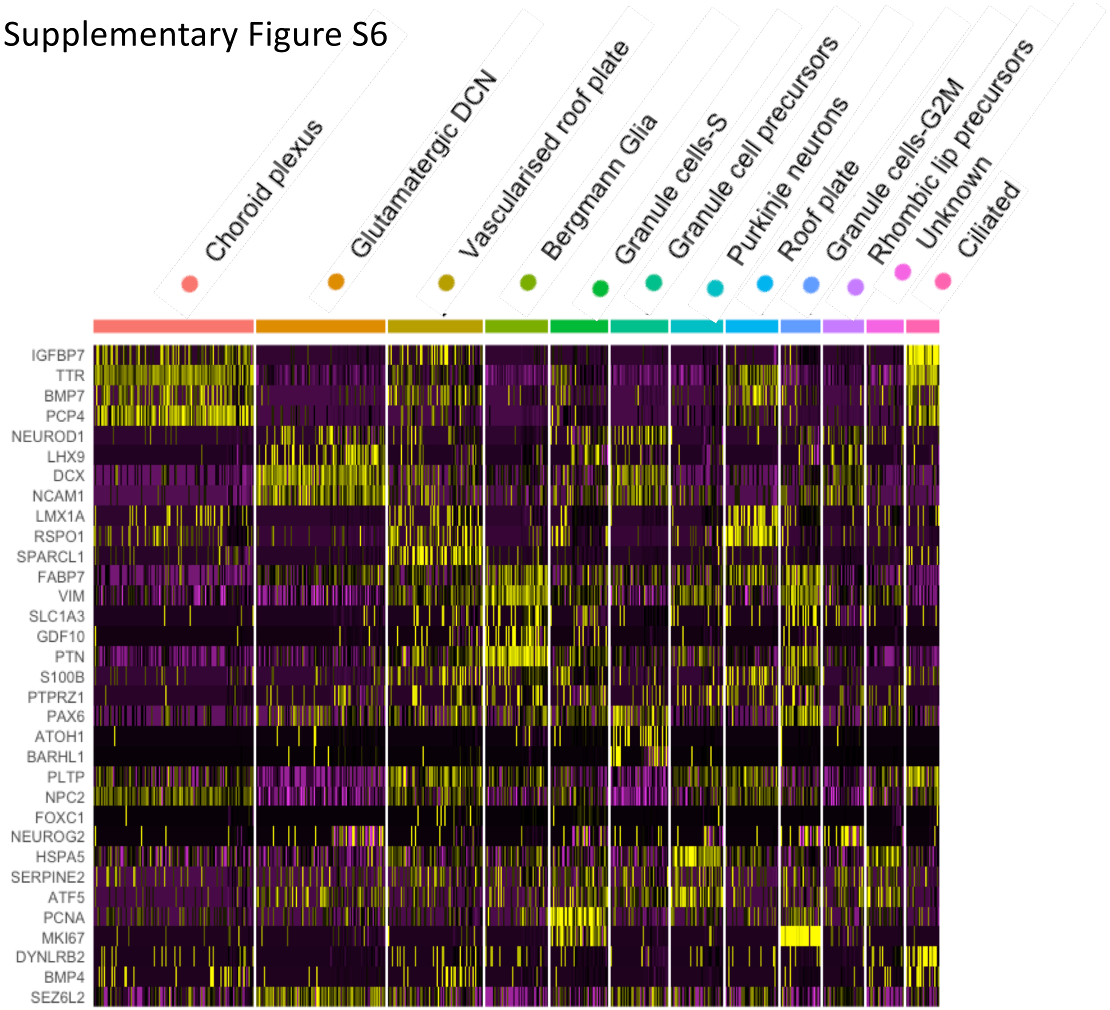


**Supplementary figure S6 –** Heatmap depicting expression of canonical cerebellar cell type-specific markers in human organoids at clustering resolution 0.8. Related to Figure 3.

### Supplementary Figure S7

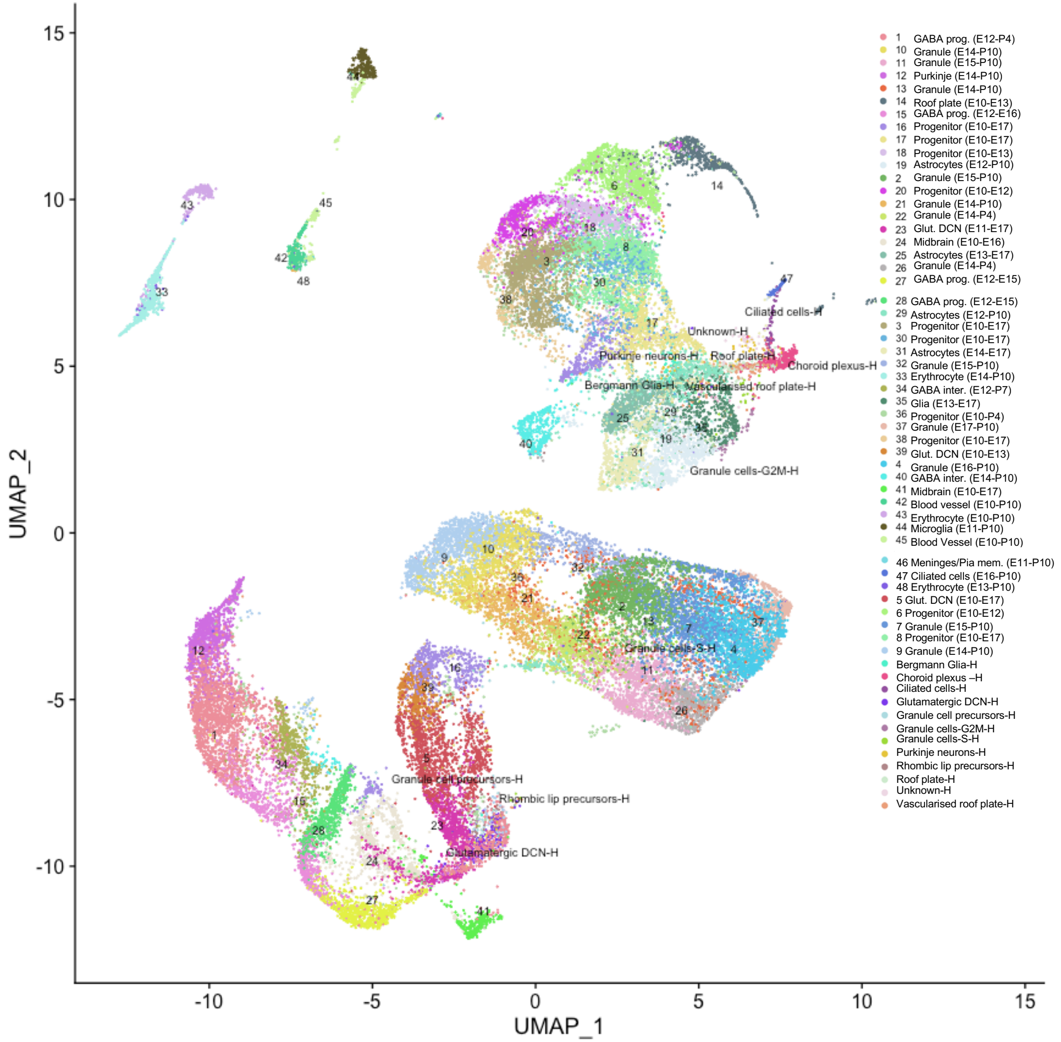


**Supplementary Figure S7 –** UMAP projection of integrated Murine and human data. Murine data is decomposed to temporally identified clusters (supplementary table 32). Related to Figure 3.

#### Supplementary Figure S8

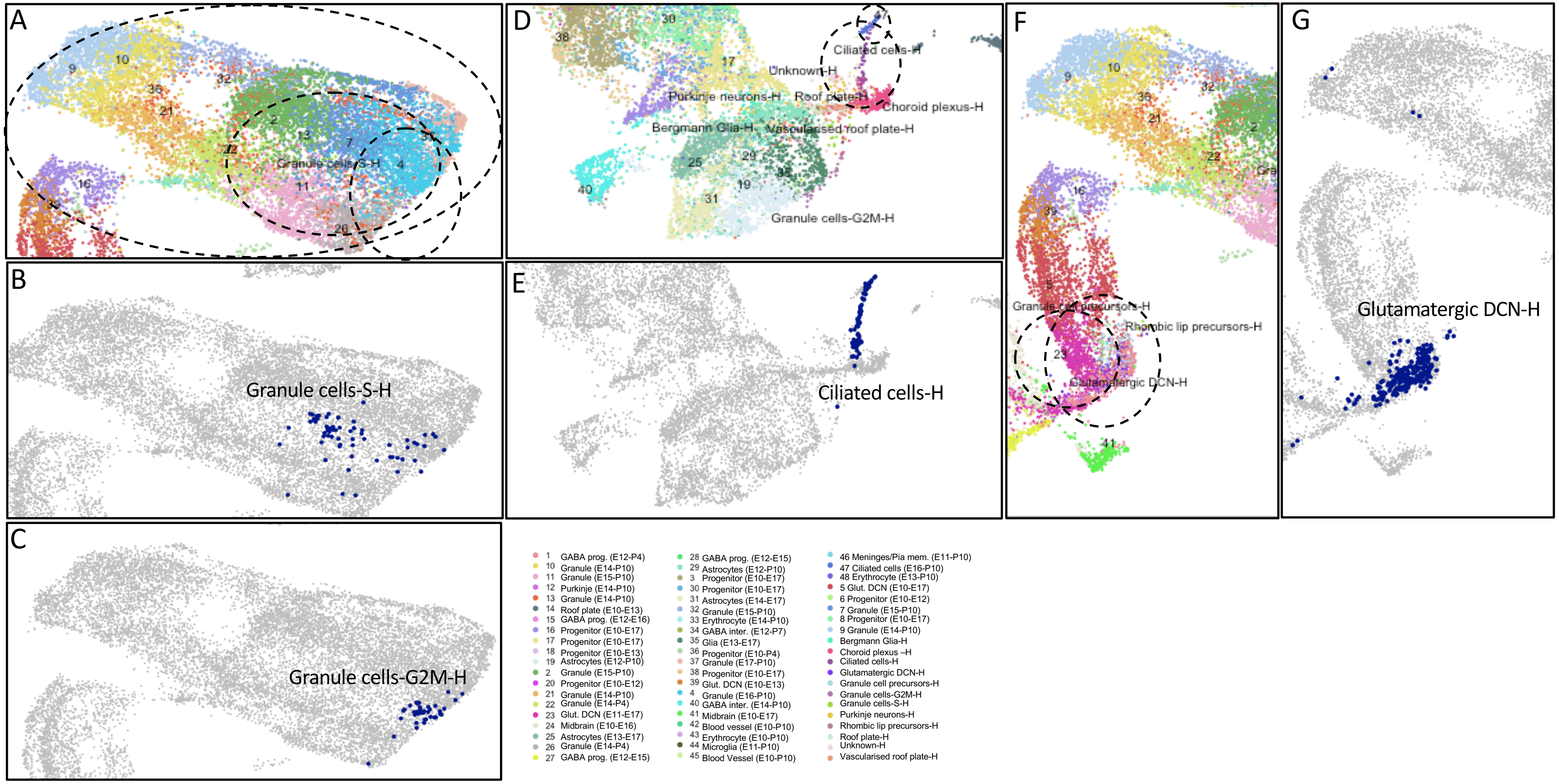


**Supplementary Figure S8 –** Select close ups of UMAP plotted according to Carter temporal metadata (Full list of clusters are described further in supplementary table 32). Related to Figure 3.

### Supplementary Figure S9

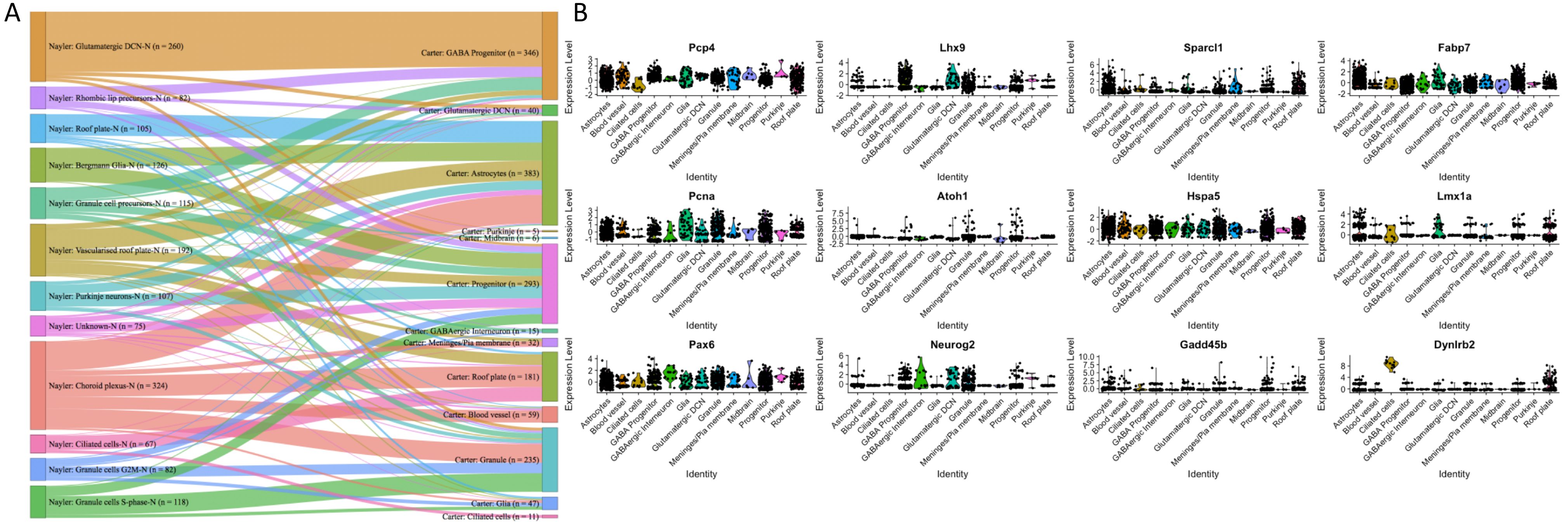


**Supplementary Figure S9 –** (A) Sankey plot of human (left) and predicted murine (right) cell types reveals high-fidelity of prediction across species following integration. (B) Violin plots reveals cluster-specific enrichment for cerebellar marker genes in human organoid data grouped according to murine predicted cell types. Related to Figure 3.

Supplementary Figure S10

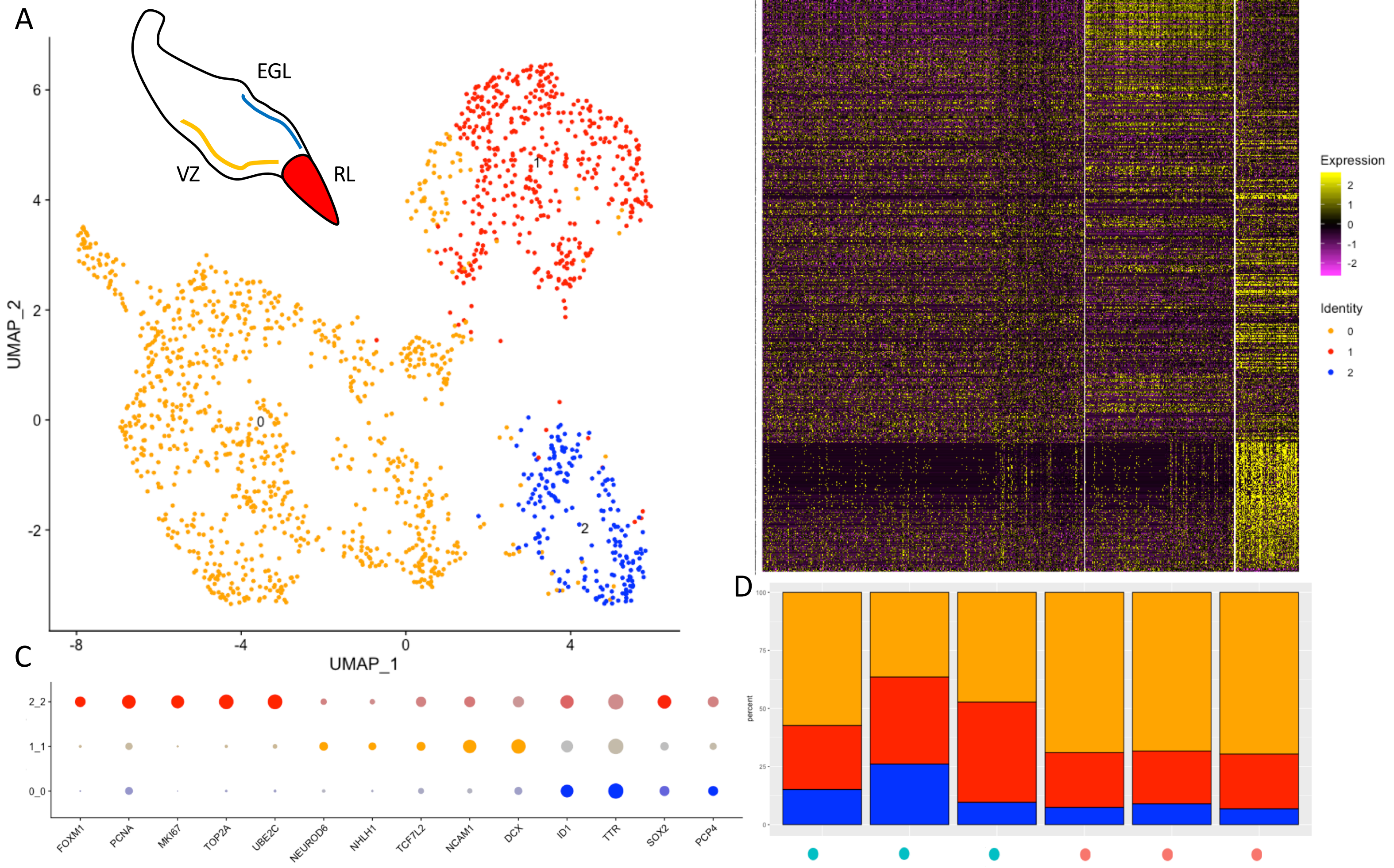


**Supplementary Figure S10 –** A) UMAP representation of clustering to resolve GABAergic/glutamatergic lineages (VZ vs RL/EGL). B) Heatmap of differentially expressed genes (supplementary table contains full list). C) Dot-plot depicting key markers driving separation of clusters. D) Bar graphs showing lineage-specification bias in Matrigel (teal) and Control (pink) organoids. Related to Figure 4.

Supplementary Figure S11

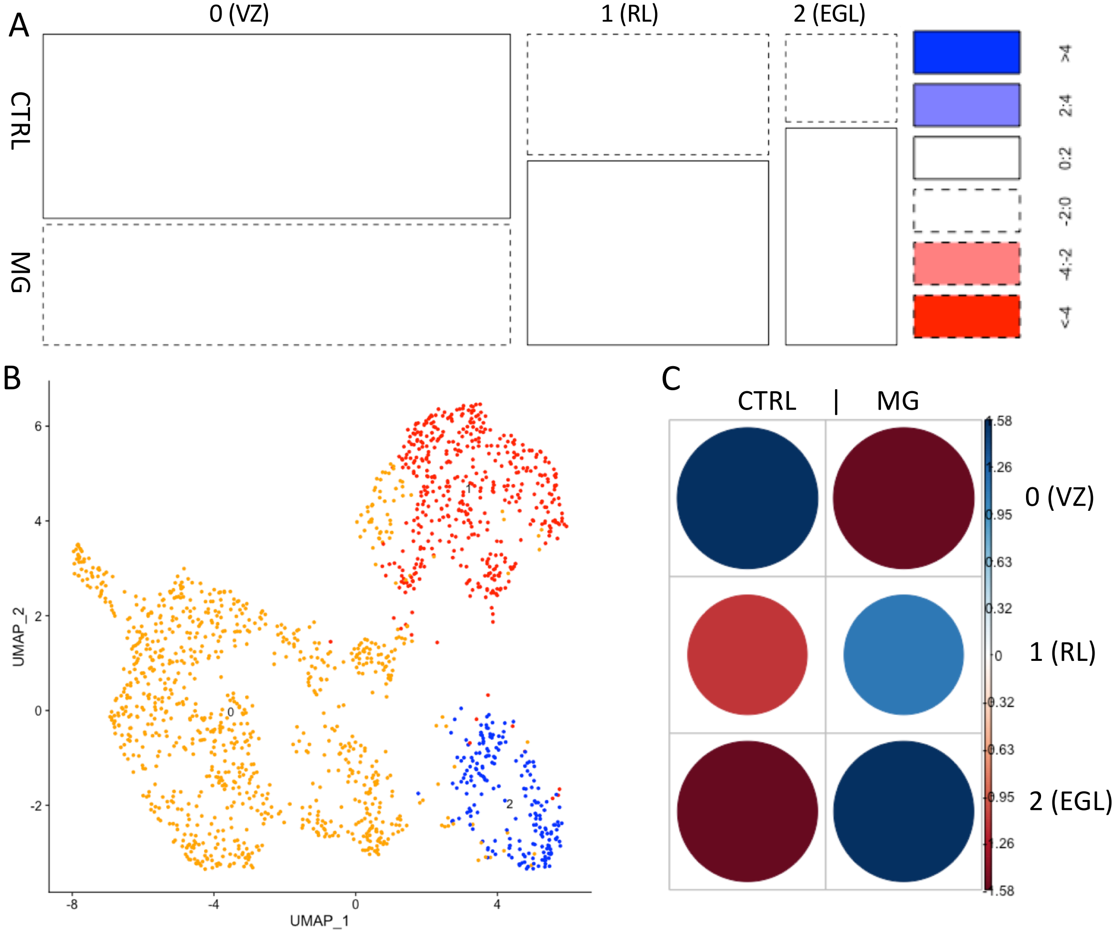


**Supplementary Figure S11 –** A) Contingency tables for chisquare tests. Standardized residuals shown for organoid population composition clustered at a resolution to resolve GABAergic/Glutamatergic lineages (VZ vs RL/EGL). B) Corresponding UMAP C) Pearson standardized residuals for corresponding clustering level. Related to Figure 4.

### Supplementary Figure S12

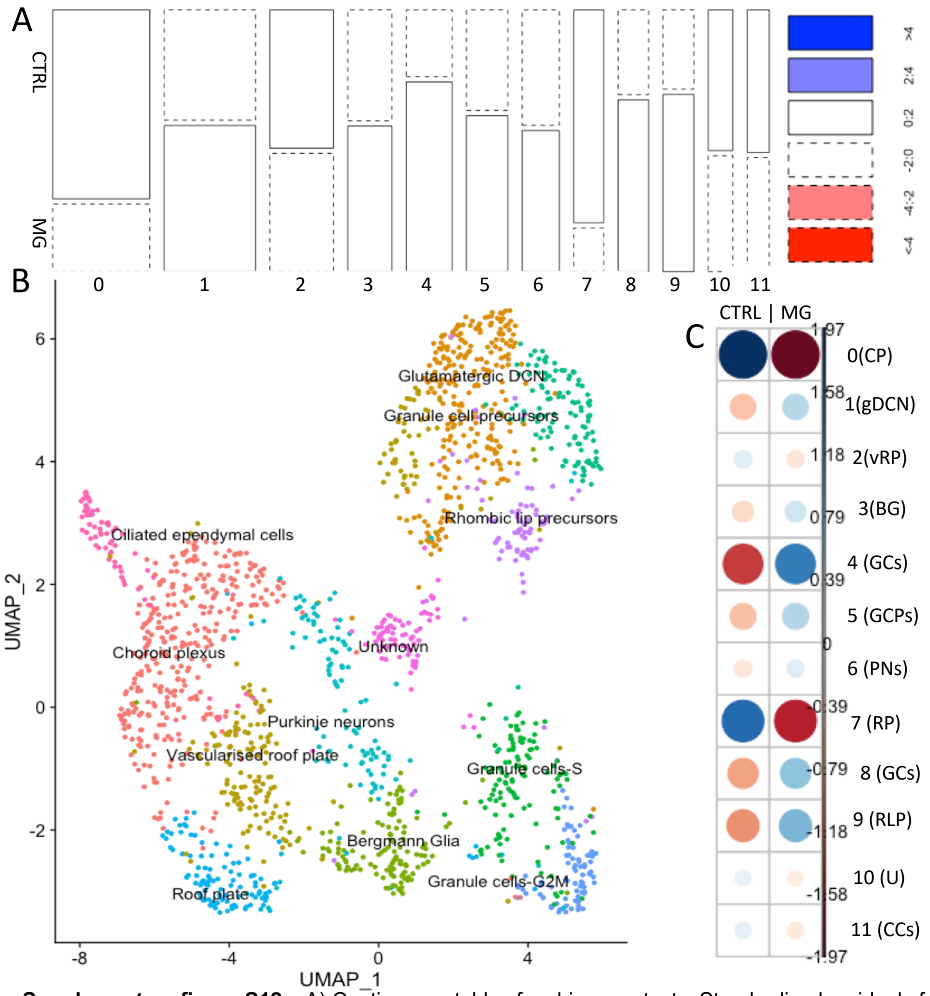


**Supplementary figure S12 –** A) Contingency tables for chisquare tests. Standardized residuals for organoid population composition clustered at a resolution to resolve cerebellar populations. B) Corresponding UMAP. C) Pearson standardized residuals for corresponding clustering level. Related to Figure 4.

#### Supplementary Figure S13

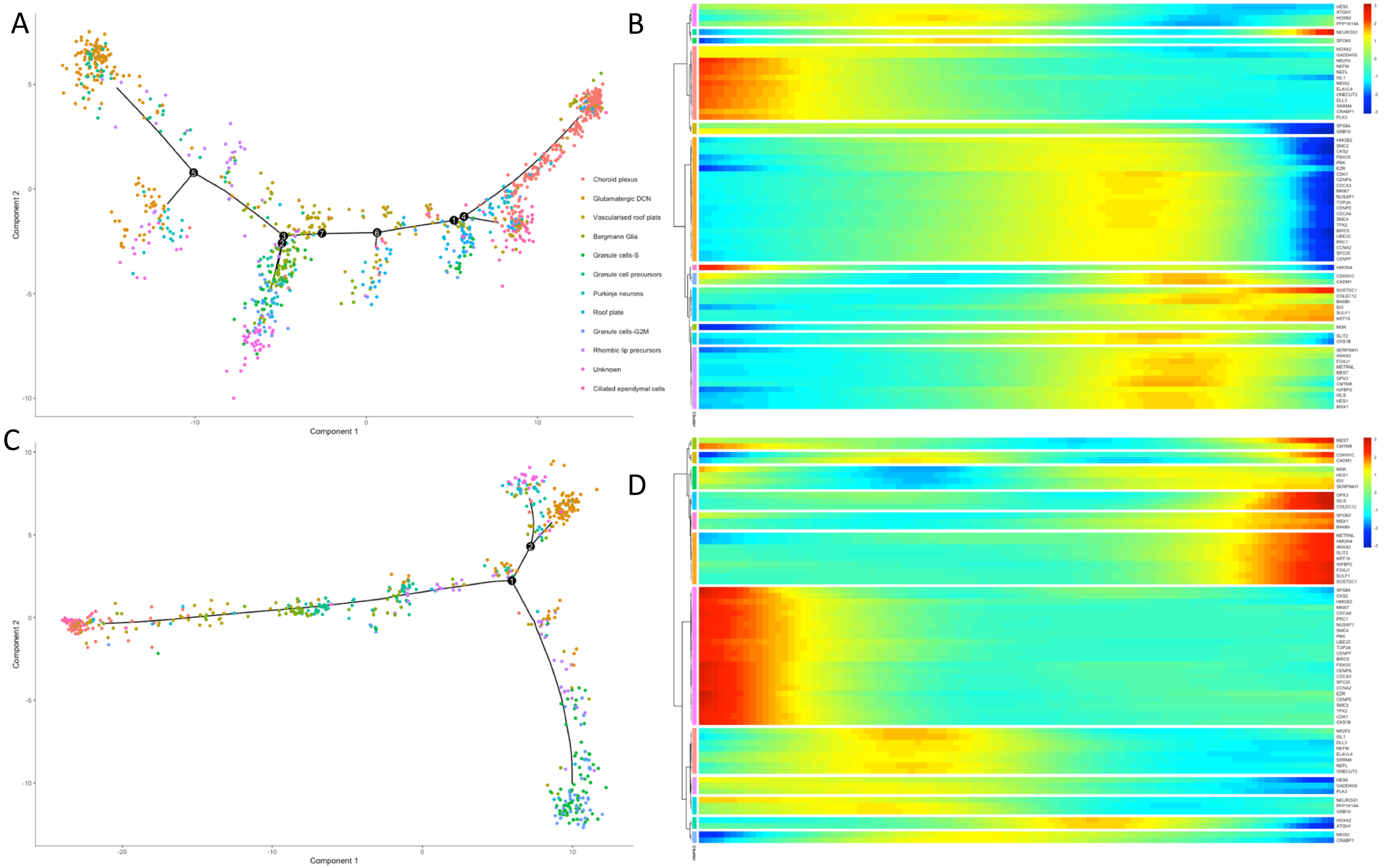


**Supplementary Figure 13 –** Pseudotime reconstructions of control (A) and Matrigel-embedded (B) organoids. Top genes driving branch separation are depicted in B and D. Related to Figure 4.
